# Multistate Enzyme Design Enables Efficient and Stereoselective Multistep Catalysis

**DOI:** 10.64898/2026.09.11.750617

**Authors:** Ngoc Thu Hang Pham, Rui Guo, Rosalinda P. Garcia Jimenez, Amy E. Hutton, Linus O. Johannissen, Johann A. Wehrstedt, Behnoush Seifinoferest, Zachary Birch-Price, Jordan Berreur, Sam Hay, Michael C. Thompson, Anthony P. Green, Roberto A. Chica

**Affiliations:** Department of Chemistry and Biomolecular Sciences, University of Ottawa, Ottawa, Ontario, K1N 6N5, Canada; Center for Catalysis Research and Innovation, University of Ottawa, Ottawa, Ontario, K1N 6N5, Canada; Department of Chemistry and Biochemistry, University of California, Merced, Merced, California 95343, United States; Manchester Institute of Biotechnology, Department of Chemistry, University of Manchester, Manchester, United Kingdom

## Abstract

Enzymes catalyze multistep reactions by stabilizing successive transition states within well organized, yet dynamic active sites. However, computational enzyme design typically targets a single transition state using rigid active-site models. Here, we introduce multistate enzyme design, which uses conformational ensembles to optimize active sites across an entire reaction coordinate. Applied to a de novo Morita–Baylis–Hillmanase, multistate enzyme design outperformed conventional single-state design, with the most active variant achieving >100-fold higher bi-substrate catalytic efficiency and surpassing an extensively optimized enzyme from directed evolution in both efficiency and enantioselectivity. Structural and kinetic analyses revealed that multistate design preserved catalytic preorganization and conformational plasticity, distributed stabilization across the reaction coordinate and avoided kinetic bottlenecks created by single-state optimization. By contrast, single-state design compromised preorganization, destabilized upstream states and shifted rate limitation away from the targeted transition state. Multistate enzyme design provides a framework for designing catalytic landscapes rather than static active sites, opening a route to efficient de novo enzymes for complex multistep chemistry.

## Introduction

Enzymes achieve remarkable catalytic efficiency by stabilizing a sequence of transition states along a reaction coordinate, often within a single preorganized active site^1–3^. For multistep reactions, this requires a delicate balance between precise positioning of catalytic residues and controlled conformational flexibility, allowing the active site to accommodate structurally distinct intermediates while maintaining productive interactions throughout the catalytic cycle^4,5^. Current computational enzyme design methods^6–9^ are poorly suited to address these requirements because they typically optimize sequences against a single transition state or reaction intermediate using largely rigid active-site models. As a result, designed enzymes for multistep reactions often show far lower turnover than their natural counterparts^10^, consistent with incomplete stabilization of the catalytic pathway. Recent attempts to mitigate this limitation through post hoc assessment of intermediate compatibility^11^ can improve activity but do not resolve the underlying issue, as they retain a single-state stabilization paradigm rather than embedding multistate optimization into the design process. Consequently, a general strategy that integrates conformational flexibility with multistate optimization across an entire reaction pathway remains lacking, constraining the design of efficient enzymes for multistep reactions with no natural precedent.

These limitations are especially apparent in efforts to design enzymes for multistep C–C bond formations. The Morita–Baylis–Hillman (MBH) reaction^12^ (Figure 1a) has emerged as a benchmark for de novo design of multistep, bi-substrate enzyme catalysis. These atom-economical transformations form a C–C bond between activated alkenes, such as α,β-unsaturated carbonyl compounds, and carbon electrophiles, such as aldehydes, via a multistep mechanism involving nucleophilic Michael addition, C–C bond formation, proton transfer and catalyst elimination. Early computational designs, such as BH32^8^, were generated using a single composite transition state and a rigid protein backbone, incorporating a catalytic histidine nucleophile and a glutamine residue intended to stabilize the enolate intermediate formed after Michael addition. Although catalytically active, BH32 required extensive directed evolution to achieve appreciable activity^13^. This process yielded BH32.12, with approximately 70-fold higher bi-substrate catalytic efficiency, driven in part by the emergence of a conformationally flexible catalytic arginine that replaced the designed glutamine and stabilized multiple oxyanion intermediates across the four-step mechanism (Figure 1a). More recently, AI-guided catalytic motif scaffolding successfully recreated the evolved His/Arg dyad of BH32.12 in de novo protein scaffolds yet still relied on single-transition-state optimization^9^, resulting in only a 3.5-fold gain in *k*_cat_ over BH32. Similarly, AI-generated NTF2-like scaffolds incorporating the His/Arg dyad efficiently reacted with a mechanistic probe reporting on early steps of the MBH mechanism, yet showed poor turnover, with conversions below those of BH32^14^. Together, these results expose a fundamental limitation of single-state enzyme design: for multistep chemistry, optimizing a single transition state does not ensure productive progression through a multistep reaction coordinate.

**Figure 1.**
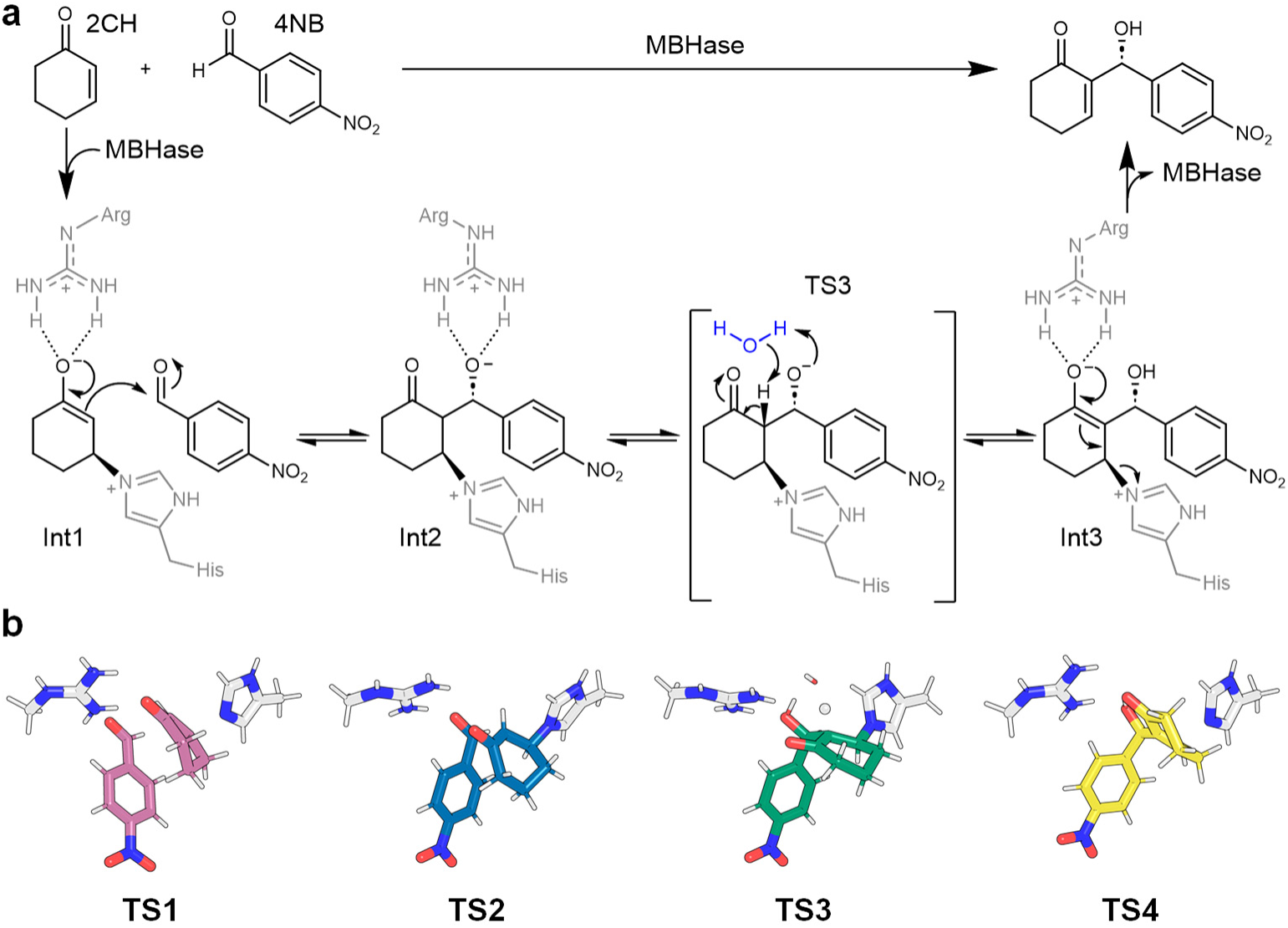
Morita–Baylis–Hillmanase mechanism. **a,** Proposed mechanism of the Morita-Baylis-Hillman reaction between 2-cyclohexen-1-one (2CH) and 4-nitrobenzaldehyde (4NB) catalyzed by BH32.12 and related MBHases. Catalysis proceeds through a four-step pathway involving a catalytic histidine nucleophile and an arginine hydrogen-bond donor that stabilizes the oxyanion intermediates (Int1-Int3) formed throughout the reaction cycle. Dotted lines indicate hydrogen bonds. TS3, which involves a water-mediated proton transfer, is the rate-limiting transition state in this proposed mechanism. **b,** Theozyme models comprising a transition state (TS, coloured) and catalytic histidine and arginine side chains (white) arranged in an optimal geometry for catalysis. The acidic proton transferred during TS3 is depicted as a sphere.

Here, we introduce multistate enzyme design, a computational approach that explicitly optimizes active sites across multiple transition states along a defined reaction coordinate. By combining distinct transition states with a backbone ensemble that captures conformational flexibility, multistate enzyme design optimizes sequence fitness across the catalytic pathway while allowing the structural rearrangements required to accommodate successive intermediates. This approach seeks to move enzyme design beyond stabilization of an individual state towards coordinated stabilization of the entire catalytic pathway.

## Results

### A computational framework for multistate optimization of enzyme reaction coordinates

To enable the design of enzymes for multistep bond-forming reactions, we developed multistate enzyme design (Figure 2). In conventional computational enzyme design^6–9,11,15,16^, a single theozyme^17^—a theoretical model of an idealized active site in which the transition state and catalytic residues are optimally arranged—is grafted onto a rigid protein scaffold to guide active-site design. By contrast, multistate enzyme design explicitly models entire reaction coordinates by combining multiple theozymes, one per transition state, with conformational ensembles of the protein backbone. In this framework, each theozyme is independently placed^18^ onto individual members of a backbone ensemble to identify protein conformations that best accommodate the required catalytic interactions. The resulting theozyme-bound protein structures then serve as templates for multistate sequence optimization^19^ to design active-site sequences predicted to stabilize the complete catalytic pathway while retaining the conformational plasticity required for multistep catalysis.

**Figure 2.**
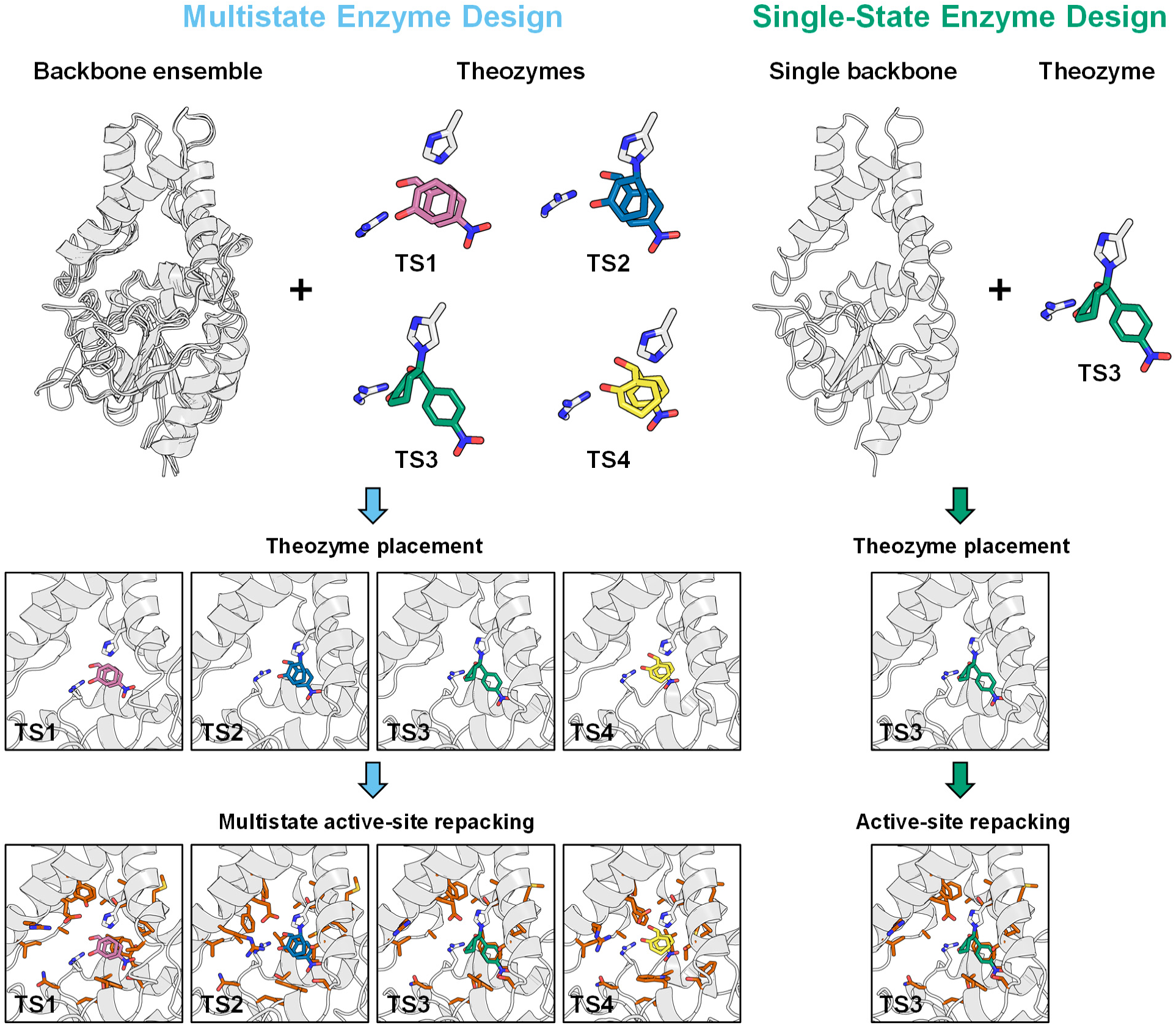
Multistate design of Morita–Baylis–Hillmanases. Multistate enzyme design takes as input a protein backbone conformational ensemble together with multiple theozymes representing distinct transition states along the reaction coordinate (TS1–TS4). Each theozyme comprises a transition state (coloured) and catalytic side chains (white) arranged in an idealized geometry derived from density functional theory calculations. Theozymes are placed onto each backbone in the ensemble to identify conformations that best reproduce the ideal geometries without steric clashes. Active-site residues (orange sticks) are then optimized by multistate design to identify sequences that stabilize all transition states. In this schematic example, the procedure yields four distinct conformations for a single protein sequence, corresponding to the modeled catalytic pathway. By contrast, single-state design uses a single protein backbone template and a single theozyme representing the rate-limiting transition state (TS3).

To implement multistate enzyme design, we used the evolved Morita–Baylis–Hillmanase BH32.12 as a model system^13^. BH32.12 catalyzes C–C bond formation between 2-cyclohexen-1-one and 4-nitrobenzaldehyde through a four-step mechanism (Figure 1a) for which high-quality theozyme models derived from density functional theory (DFT) calculations are available for each catalytic step (TS1–TS4, Figure 1b). Because the catalytic geometries differ across the reaction coordinate, we hypothesized that accurate accommodation of all four theozymes would require explicit treatment of backbone flexibility. To capture this flexibility, we generated conformational ensembles for BH32.12 and its evolutionary precursor BH32.8 by crystallographic ensemble refinement^20^ (Methods, Supplementary Table 1), a method that models conformational heterogeneity by representing experimental electron density as an ensemble of structurally distinct protein conformations. We previously showed that this approach can reveal conformational substates that accommodate transition-state geometries and catalytic interactions more accurately than the single average crystal structure^21^, thereby enabling the design of more catalytically productive active sites^22,23^.

Using the physics-based protein design software Triad^21,24^, we independently placed each of the four theozymes onto every backbone in the BH32.12 and BH32.8 ensembles while preserving the DFT-derived interactions with catalytic residues His23 and Arg124 (Supplementary Tables 2–3). All four theozymes could be incorporated with high-fidelity on distinct conformations from the ensembles without steric clashes. Notably, neither the average crystal structures nor any single ensemble member could simultaneously support accurate placement of all four theozymes. These findings indicate that stabilization of the full reaction coordinate requires an ensemble-based representation of active-site conformational space.

We next selected the four theozyme-bound structures displaying the lowest RMSD relative to the DFT models, one per TS (Supplementary Figure 1), as templates for multistate sequence design in Triad. In multistate design^25^, every state, in this case the four backbone conformations with distinct placed theozyme, are used to guide the sequence search. We implemented a multistate fitness function^19^ that prioritized stabilization of the rate-limiting transition state, TS3^13^, while penalizing destabilization of the remaining states relative to the BH32.12 sequence (Methods). In parallel, we performed conventional single-state design calculations using only the TS3-bound structure to generate sequences optimized exclusively for stabilization of the rate-limiting transition state^13^.

Both design strategies searched the same sequence space comprising hydrophobic and hydrogen-bonding amino acids predicted to support substrate recognition (Supplementary Table 4). We generated 1,000 sequences from each approach and filtered them based on low TS3 energy and minimal energetic variation across the reaction coordinate; the latter criterion was not applied during single-state design because only a single transition state was considered. Predicted energy profiles (Supplementary Figure 2a) showed that multistate design sequences maintained favorable energies across the entire reaction coordinate, with only modest destabilization of TS2 relative to BH32.12. By contrast, single-state designs stabilized TS3 more efficiently but greatly destabilized one or more additional transition states, highlighting a central limitation of conventional single-transition-state optimization. From these designs, ten multistate variants (M1–10) containing 8–10 active-site mutations relative to BH32.12 and ten single-state variants (S1–10) containing 12–13 mutations were selected for experimental characterization (Supplementary Table 5).

### Multistate design yields more active and selective enzymes than single-state design

All variants generated by multistate design and single-state optimization were soluble, folded and stable (Supplementary Table 6, Supplementary Figure 3–4). Activity assays revealed that all but one multistate design were significantly more active than the single-state designs (Figure 3a). Six multistate variants exceeded the activity of BH32.8, whereas none of the single-state designs achieved half the conversion of BH32.8. The most active multistate variant, BH32.M6, contains nine active-site mutations (Figure 3b, Supplementary Table 5, Supplementary Figure 5), matched the conversion of BH32.12 and achieved up to 44-fold higher conversion than the single-state designs. These results indicate that optimization of a single rate-limiting transition state, even in the presence of the catalytic His23/Arg124 motif, is insufficient to achieve efficient catalysis of this multistep reaction.

**Figure 3.**
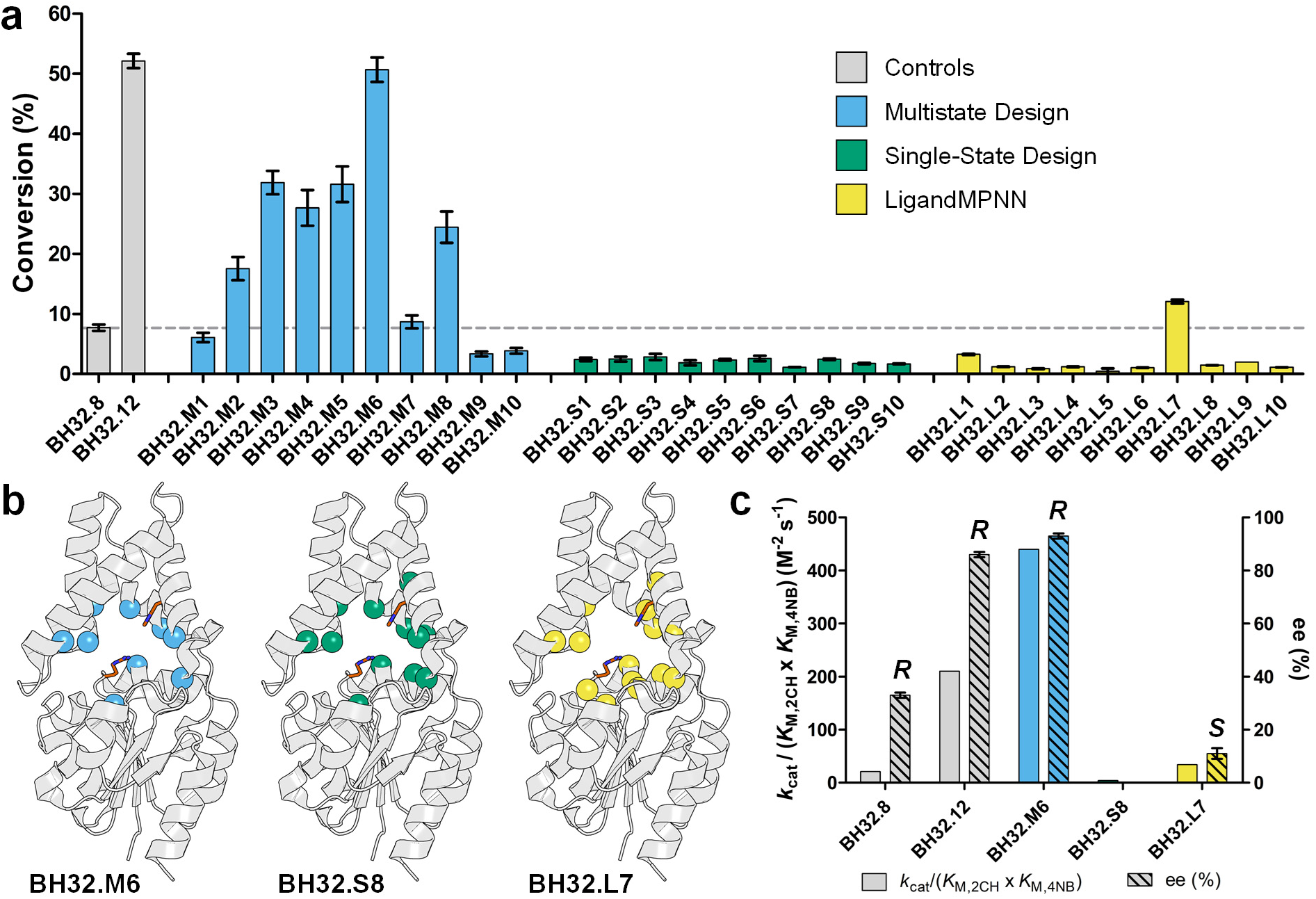
Activity of designed MBHases. **a,** Product conversion for designed variants and control MBHases (mean ± s.d., n = 2–14). The dashed line indicates the conversion achieved by BH32.8. Enzyme reactions were performed with 3 mM 2-cyclohexen-1-one, 0.6 mM 4-nitrobenzaldehyde, and 20 µM enzyme (3.3 mol%) at 30 °C in phosphate-buffered saline (pH 7.4) containing 3% (v/v) acetonitrile, and analyzed after 22 h. **b,** Structural representation of mutated residues (spheres) mapped onto the crystal structure of BH32.12 (PDB ID: 6Z1L). BH32.M6, BH32.S8, and BH32.L7 contain 9, 12, or 14 active-site mutations relative to BH32.12. Catalytic residues His23 and Arg124 are shown as orange sticks. **c,** Bi-substrate catalytic efficiency and enantiomeric excess for MBHase variants (see Table 1 for data). 2CH and 4NB denote 2-cyclohexen-1-one and 4-nitrobenzaldehyde, respectively. The enantioselectivity of BH32.S8 could not be determined because the aldol by-product interfered with reliable quantification of the product enantiomers. All other enzymes predominantly produce the *R* enantiomer of the product, except BH32.L7, which favors the *S* enantiomer.

**Table 1.** Kinetic parameters of various MBHases.

| Enzyme | $k_{\text{cat}}$<br>($\text{min}^{-1}$) | $K_{\text{M},2\text{CH}}$<br>($\text{mM}$) | $K_{\text{M},4\text{NB}}$<br>($\text{mM}$) | $k_{\text{cat}}/K_{\text{M},2\text{CH}}$<br>($\text{M}^{-1} \text{s}^{-1}$) | $k_{\text{cat}}/K_{\text{M},4\text{NB}}$<br>($\text{M}^{-1} \text{s}^{-1}$) | $k_{\text{cat}}/(K_{\text{M},2\text{CH}} \times K_{\text{M},4\text{NB}})$<br>($\text{M}^{-2} \text{s}^{-1}$) | ee (R)<br>(%) |
| --- | --- | --- | --- | --- | --- | --- | --- |
| BH32.8 | $0.0109 \pm 0.0007$ | $4.9 \pm 0.4$ | $1.8 \pm 0.2$ | 0.037 | 0.10 | 21 | $33 \pm 1$ |
| BH32.12 | $0.058 \pm 0.003$ | $4.3 \pm 0.3$ | $1.05 \pm 0.09$ | 0.23 | 0.92 | 210 | $86 \pm 1$ |
| BH32.M6 | $0.031 \pm 0.006$ | $0.84 \pm 0.02$ | $1.4 \pm 0.5$ | 0.62 | 0.37 | 440 | $93 \pm 1$ |
| BH32.S8 | $0.0021 \pm 0.0002$ | $8 \pm 1$ | $1.0 \pm 0.1$ | 0.004 | 0.035 | 4 | N.D. |
| BH32.L7 | $0.0100 \pm 0.0005$ | $2.7 \pm 0.2$ | $1.8 \pm 0.2$ | 0.062 | 0.093 | 34 | $-11 \pm 2$ |

To determine whether the enhanced activity of BH32.M6 arose from improved turnover, productive substrate binding, or both, we kinetically characterized BH32.M6 alongside one of the most active single-state designs, BH32.S8, which contains 12 mutations (Figure 3b, Supplementary Table 5, Supplementary Figure 5), and compared both variants with BH32.8 and BH32.12. BH32.M6 displayed a 15-fold higher *k*_cat_ than BH32.S8 together with a 10-fold lower *K*_M_ for 2-cyclohexen-1-one, while its *K*_M_ for 4-nitrobenzaldehyde did not vary significantly (Table 1, Supplementary Figure 6). As a result, BH32.M6 exhibited a >100-fold increase in bi-substrate catalytic efficiency relative to BH32.S8 (Figure 3c). Moreover, while BH32.M6 produced the target MBH adduct with high chemoselectivity, BH32.S8 produced a significant proportion of a competing aldol by-product (Supplementary Figure 7). BH32.M6 was also twofold more catalytically efficient than BH32.12 mostly due to improved productive binding for 2-cyclohexen-1-one. BH32.M6 was also more stereoselective, achieving 93% ee(R), compared with 86% for BH32.12 (Figure 3c, Supplementary Figure 8). Together, these results demonstrate that multistate enzyme design can produce biocatalysts that are both more catalytically efficient and more stereoselective than conventional single-state enzyme design.

### Deep-learning sequence design yields limited catalytic improvement

Having established that multistate design outperforms single-state optimization, we next asked whether a deep learning-based approach could achieve similar gains. We adapted LigandMPNN^26^ for multiconformer design^27^ by providing the four theozyme-bound structures as a homotetramer and restricting sequence optimization to the same active-site positions sampled in our physics-based multistate calculations. We generated 100 sequences and selected ten whose AlphaFold2^28^ models yielded the highest predicted structural confidence, the closest agreement with the BH32.12 crystal structure, and accurate positioning of the catalytic residues (Methods). To maximize sequence diversity, we additionally required that each selected design differ from the others by at least two active-site mutations. This process yielded ten LigandMPNN variants (L1–10), containing 12–14 mutations relative to BH32.12 (Supplementary Table 5).

All ten LigandMPNN variants were soluble, folded and stable (Supplementary Table 6, Supplementary Figures 3–4). However, catalytic gains were limited: only BH32.L7 exceeded the activity of BH32.8, whereas the other variants showed conversions comparable to the single-state designs (Figure 3a). BH32.L7 (Figure 3b, Supplementary Table 5, Supplementary Figure 5) achieved approximately fourfold lower conversion than the best physics-based multistate design, BH32.M6. Its *k*_cat_ was fivefold higher than BH32.S8 but remained threefold and sixfold lower than those of BH32.M6 and BH32.12, respectively (Table 1, Supplementary Figure 6). Although BH32.L7 displayed improved *K*_M_ for 2-cyclohexen-1-one relative to BH32.12, its overall bi-substrate catalytic efficiency remained an order of magnitude below that of BH32.M6 (Figure 3c). Notably, BH32.L7 also exhibited poor and inverted stereoselectivity (Supplementary Figure 8), producing the opposite enantiomer with −11% ee(R), despite the correct stereochemical configuration being specified in the input design models.

We sought to determine whether differences in transition-state energetics could explain these limitations. Triad evaluation of the LigandMPNN designs yielded energy profiles broadly compatible with accommodation of all four transition states, but with substantially greater energetic imbalance than the physics-based multistate designs (Supplementary Figure 2b). In particular, the profiles preferentially stabilized TS2 while destabilizing TS3. Thus, in this implementation, LigandMPNN produced well-folded variants capable of accommodating multiple transition-state geometries, but did not achieve the energetic balance, high catalytic efficiency and stereochemical control obtained with physics-based multistate optimization. These results highlight the benefit of explicitly incorporating transition-state energetics and stereochemical constraints into sequence optimization for this catalytic system.

### Multistate design preorganizes catalytic residues without overpacking the active site

To uncover the structural basis of the enhanced activity and selectivity of BH32.M6, we solved noncryogenic crystal structures of BH32.M6, BH32.S8 and BH32.L7 in their unbound state (Supplementary Tables 7–8). We chose noncryogenic crystallography because it can capture conformational heterogeneity that is often obscured at cryogenic temperatures, providing a more accurate view of the structural ensemble sampled by the protein^29,30^. All three variants closely resembled the parental BH32.12 scaffold (Figure 4a), except for residues 184–188, which are disordered in BH32.12 but adopt a π-helix in BH32.L7 and a helix–turn–bend–helix motif in BH32.M6 and BH32.S8 (Figure 4b). Despite their similar backbone structures, BH32.S8 showed pronounced deviations in catalytic-residue conformations, whereas BH32.M6 and BH32.L7 retained conformations similar to those of BH32.12 (Figure 4c), suggesting a fundamentally distinct active-site architecture in the single-state variant. Cavity-volume analysis^31^ confirmed this distinction: BH32.S8 formed a more compact active site than BH32.M6, BH32.L7 and BH32.12 (Supplementary Figure 9). Consistent with the computed energy profiles (Supplementary Figure 2c), the tighter active site of BH32.S8 likely favours stabilization of TS3 but comes at the expense of other transition states. This finding highlights a central limitation of conventional single-transition-state optimization, which favours tight packing around a single transition state without accounting for the structural requirements of other states along the reaction coordinate.

**Figure 4.**
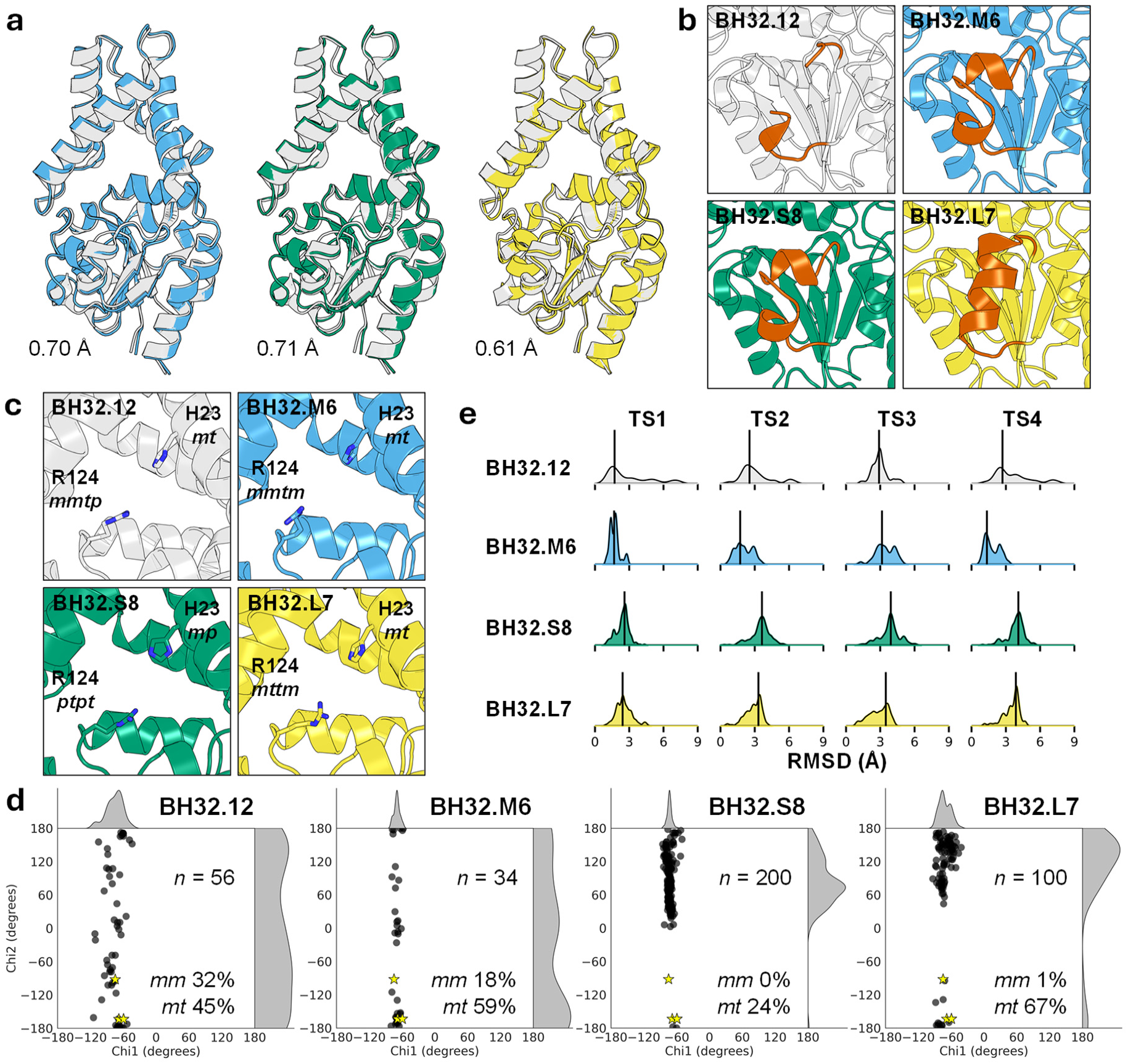
Crystal structures. BH32.12, BH32.M6, BH32.S8 and BH32.L7 are colored white, blue, green and yellow, respectively. **a,** Structural overlay showing that the designed active-site mutations do not substantially alter the overall protein fold. Backbone root-mean-square deviations (RMSDs) relative to BH32.12 are indicated. **b,** The BH32.12 structure lacks electron density for residues 184–188, indicating conformational heterogeneity in this region. In BH32.M6 and BH32.S8, residues 181–193 (orange) adopt a helix–turn–bend–helix motif, whereas BH32.L7 contains a π-helix. **c,** His/Arg catalytic dyad. Rotamer nomenclature follows the convention in which *m* denotes a *gauche^−^* conformation (χ angle in the 0 to −120° rotameric well), *p* denotes a *gauche^+^* conformation (χ angle in the 0 to +120° well), and *t* denotes a *trans* conformation (χ angle in the +120 to −120° well). Dihedral angles are listed in the order χ_1_, χ_2_, χ_3_, and χ_4_. **d,** χ_1_–χ_2_ distributions for catalytic His23 derived from ensemble refinement; the number of conformers in each ensemble (*n*) is indicated. BH32.12 and BH32.M6 sample a broader set of rotamers (black dots), overlapping with those observed in the TS1–TS4 design models (yellow stars). Percentages indicate the fraction of ensemble structures sampling these conformations. BH32.S8 does not sample the *mm* rotamer and samples *mt* rotamers at lower frequency and with larger deviations in χ_2_ relative to the design models. BH32.L7 samples *mm* rotamers at substantially lower frequency than BH32.12 and BH32.M6. **e,** Kernel density estimation distributions of RMSDs between the guanidinium group of catalytic Arg124 in each ensemble conformer and the four corresponding theozyme-bound structures (TS1–TS4). The black line indicates the mode of each distribution.

We next evaluated whether multistate enzyme design improves catalytic preorganization, which is the precise positioning of catalytic residues in the absence of substrate, a key determinant of catalytic efficiency. Ensemble refinement of the crystallographic data^20^ (Supplementary Table 1) revealed pronounced differences in active-site conformational heterogeneity. For the catalytic His23 residue, BH32.12 and BH32.M6 sampled a broad range of rotamers, including those represented in the TS1–TS4 design models, in 77% of structures in their respective ensemble (Figure 4d), despite BH32.12 being solved at cryogenic temperature. By contrast, BH32.S8 sampled a narrower range of His23 conformations, with productive rotamers observed in only 24% of ensemble structures and showing greater χ_2_ deviations from the design models. BH32.L7 showed an intermediate phenotype, sampling the productive rotamers in 68% of structures.

For the other catalytic residue, Arg124, we instead calculated the RMSD of its guanidinium group relative to each of the four superposed theozyme models since its four rotatable side-chain bonds preclude straightforward χ_1_–χ_2_ analysis. BH32.M6 exhibited lower modal RMSDs for three of the four transition states than all other variants, including BH32.12 (Figure 4e), indicating that the most highly populated conformations of Arg124 more closely resembled the corresponding theozyme geometries. For TS3, however, the modal RMSD was slightly higher than that of BH32.12. Notably, BH32.S8 exhibited the highest modal RMSD across all four transition states despite relatively narrow distributions, indicating that Arg124 adopts a well-defined but less catalytically preorganized conformation. Together, these structures reveal a clear distinction between the two design strategies: multistate enzyme design maintains an active-site conformational ensemble that is well preorganized for multiple transition states, whereas single-state design produces a more constrained active site with reduced compatibility across the reaction coordinate.

### Multistate enzyme design balances catalytic barriers across the reaction coordinate

We next investigated whether multistate enzyme design distributes catalytic barriers more evenly across the reaction coordinate than conventional single-state design, given its ability to produce active sites for stabilizing multiple transition states. We first tested whether TS3 remained rate-limiting using kinetic isotope effects (KIEs) with deuterated 2-cyclohexen-1-one^32^. Like BH32.12, BH32.M6 showed a significant KIE (Figure 5a), consistent with rate-limiting proton transfer at TS3^13^. By contrast, BH32.S8 showed no detectable KIE, indicating that proton transfer at TS3 no longer contributes substantially to rate limitation and implicating another step in the catalytic cycle.

**Figure 5.**
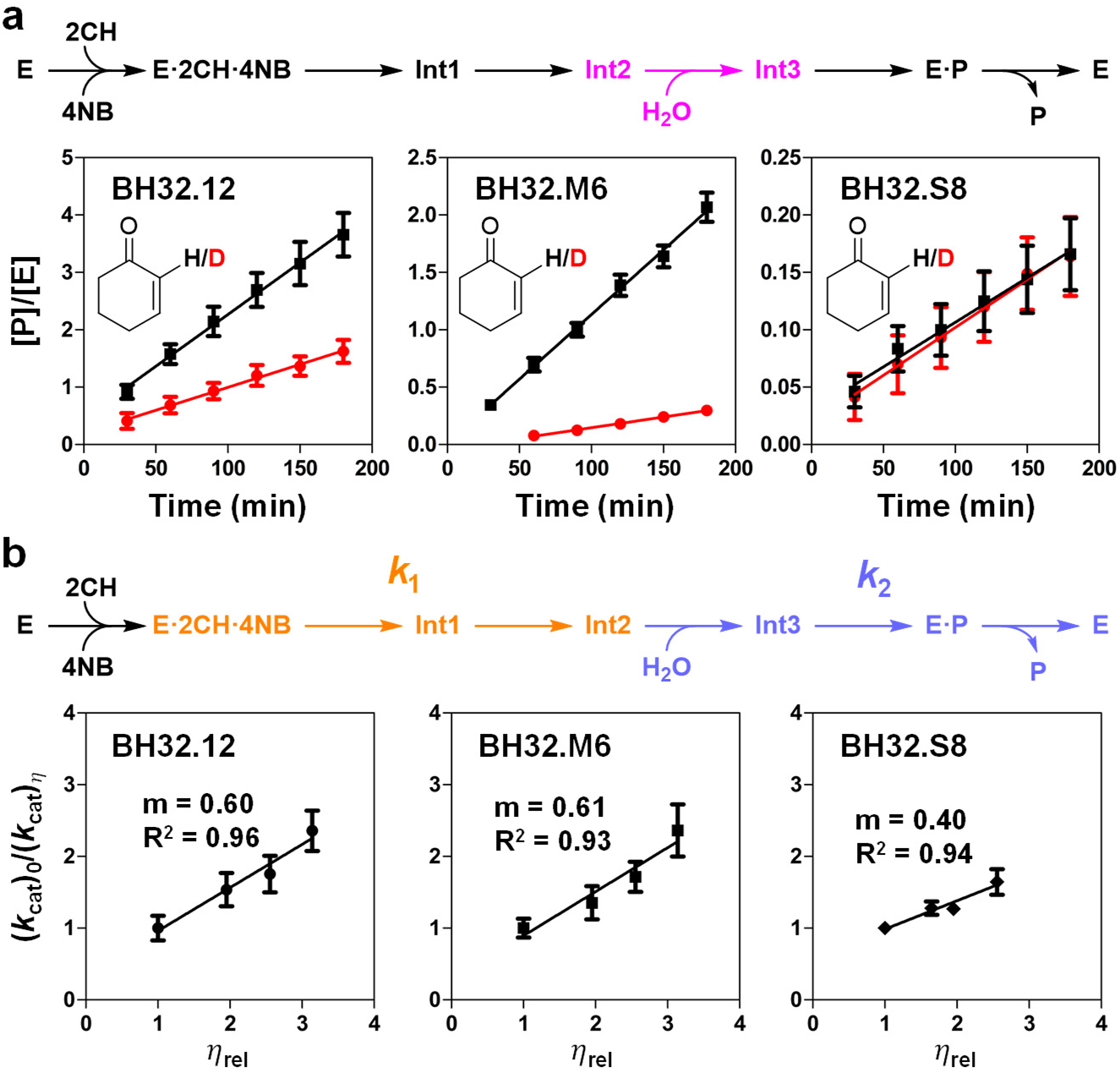
Mechanistic investigation of MBHase catalysis. **a,** Reaction scheme for the Morita–Baylis– Hillman reaction. E, 2CH, 4NB, Int1–3 and P denote enzyme, 2-cyclohexen-1-one, 4-nitrobenzaldehyde, intermediates 1–3 and product, respectively. Kinetic isotope effect (KIE) plots comparing reaction rates with 2-cyclohexen-1-one (black) or 2-deuterocyclohex-2-en-1-one (red) across different MBHases. The mechanistic step probed by the KIE, corresponding to TS3, is highlighted in magenta. Reactions were performed with 2CH (10 mM) and 4NB (2 mM) in phosphate-buffered saline (pH 7.4) containing 3% (v/v) acetonitrile as cosolvent. Data represent mean ± s.e.m. from n = 2 independent protein batches for BH32.M6 or n = 3 three independent protein batches for BH32.12 and BH32.S8, with one technical replicate per batch. **b,** Kinetic solvent viscosity effects on *k*_cat_. *k*_cat_ values measured at different relative viscosities (*η*_rel_) were normalized to values measured in non-viscous buffer and plotted against *η*_rel_, revealing a linear relationship. Slopes between 0 and 1 indicate that turnover is partially limited by both diffusion-dependent processes (*k*_2_, blue) and diffusion-independent chemical steps (*k*_1_, orange), with slopes (m) > 0.5 indicating *k*_1_ > *k*_2_ and m < 0.5 indicating *k*_1_ < *k*_2_. Data represent mean ± s.e.m. from n = 3–6 independent biological replicates measured at different buffer viscosities.

To identify the new bottleneck in BH32.S8, we measured kinetic solvent viscosity effects (KSVEs) for 2-cyclohexen-1-one using sucrose as the viscogen. KSVEs on *k*_cat_ distinguish diffusion-dependent processes contributing to turnover, such as product release and conformational rearrangements, from diffusion-independent chemical steps occurring within the active site, which is expected to be shielded from bulk solvent effects after formation of the Michaelis complex^33^. In the proposed MBHase mechanism (Figure 1a), water entry at TS3 and product release are expected to be diffusion-dependent, whereas the preceding chemistry through TS1 and TS2 is diffusion-independent. We therefore divided the catalytic cycle into two segments (Figure 5b): from the Michaelis complex to Int2, represented by the diffusion-independent rate constant *k*_1_, and from Int2 to product release, represented by the diffusion-dependent rate constant *k*_2_.

BH32.M6 showed a KSVE slope nearly identical to that of BH32.12 and greater than 0.5 (Figure 5b), indicating that turnover is partially limited by both downstream diffusion-dependent processes and upstream chemical steps, with *k*_2_ being slower than *k*_1_. This kinetic partitioning is consistent with rate-limiting proton transfer at TS3 for both enzymes (Figure 5a). By contrast, BH32.S8 showed a slope below 0.5, indicating within this kinetic framework that the upstream chemical steps had become slower than the downstream diffusion-dependent steps. Consistent with this interpretation, random-acceleration molecular dynamics simulations^34^ of Triad-generated product-bound complexes showed greater product-release propensity for BH32.S8 than for BH32.M6 and BH32.12 (Supplementary Figure 10), suggesting that product release is unlikely to be rate-limiting in the single-state variant. Together with the computed destabilization of TS1 and TS2 for BH32.S8 (Supplementary Figure 2c) and the absence of a KIE, these results suggest that rate limitation has moved from TS3 to an upstream chemical step.

To probe the early catalytic steps, we used a mechanistic inhibitor that undergoes Michael addition with the catalytic histidine to irreversibly trap an Int2 analogue (Supplementary Figure 11a)^13^. The inactivation rate (*k*_inact_) therefore provides a measure of how rapidly the initial Michael addition occurs following formation of the enzyme–inhibitor complex. BH32.12 was inactivated too rapidly to yield reliable inhibition constants (Supplementary Figure 11b), but BH32.M6 showed fourfold higher *k*_inact_ than BH32.S8 (Supplementary Table 9, Supplementary Figure 11c). This result suggests that the multistate variant undergoes Michael addition more rapidly than the single-state variant, consistent with slower passage through TS1 in BH32.S8. Together with the KSVE and KIE measurements, and computed energy profiles (Supplementary Figure 2c), these data support the presence of an upstream barrier in BH32.S8.

Overall, these mechanistic investigations reveal distinct kinetic consequences of the two design strategies. Multistate enzyme design preserves a balanced catalytic landscape: BH32.M6 retains the parental rate-limiting step at TS3 without introducing a comparably large upstream barrier. Single-state optimization instead redistributes the catalytic barriers, favouring TS3 stabilization at the expense of upstream states and shifting rate limitation to an earlier step. Efficient multistep catalysis therefore requires not simply lowering the highest barrier but coordinating stabilization across the entire reaction coordinate.

## Discussion

De novo design of efficient multistep, bond-forming enzymes poses a fundamental challenge: multiple, distinct catalytic states impose competing structural and energetic requirements that must be accommodated within a single active site^1,4^. Our results show that these requirements can be programmed directly through multistate enzyme design. By scaffolding multiple theozymes across experimentally resolved protein conformational ensembles to design active sites, multistate enzyme design produced substantially more catalytically efficient and stereoselective enzymes than conventional single-state design. Rather than maximizing complementarity to one transition state, multistate enzyme design maintained productive configurations across the reaction coordinate. Single-state optimization instead stabilized the target transition state at the expense of alternate states, redistributing barriers and creating new kinetic bottlenecks. These findings provide direct experimental evidence that efficient multistep catalysis requires coordinated stabilization across multiple catalytic states along the reaction coordinate, challenging the prevailing paradigm in de novo enzyme design of optimizing sequences for individual transition states^6–9,11,15^.

This limitation of single-state design may be general. For example, the bi-substrate catalytic efficiency of BH32.S8 (4 M^-1^ s^-1^) is comparable to that of the most active MBHase designed on a de novo protein scaffold using the same His/Arg catalytic dyad (2 M^-1^ s^-1^)^9^. Although these enzymes were designed against a different transition state (TS3 here versus TS1 in ^9^), this convergence points to a fundamental limitation of single-state optimization: improving complementarity to one catalytic state does not necessarily increase efficiency when other states remain unstabilized. Consistent with our findings, recent de novo design of serine hydrolases^11^ showed that filtering designs for compatibility with multiple reaction intermediates yielded more efficient artificial enzymes. In that approach, however, multistate compatibility was primarily used as a post-design filter. Our multistate enzyme design method instead incorporates multiple transition states directly into sequence optimization, allowing their competing geometric and energetic requirements to be balanced during the search. This distinction is important: post hoc filtering can identify single-state designs that happen to accommodate additional states but cannot recover sequences excluded by the single-state design objective. Multistate enzyme design therefore shifts multistate compatibility from a property to be selected after design to a constraint that shapes the design search itself.

This advance was enabled by our previous development of ensemble-based enzyme design^21^, which has shown that crystallographic ensembles can reveal protein conformations that better accommodate individual theozymes and thereby improve catalytic activity^22,23^. Multistate enzyme design extends this principle from individual transition states to the reaction coordinate. No single conformation could accurately accommodate all four theozymes, demonstrating that active-site heterogeneity is not incidental but integral to catalysis. Multistate enzyme design exploits this heterogeneity by allowing distinct catalytic states to be accommodated by different, interconverting conformations while preserving catalytic preorganization. The resulting BH32.M6 enzyme retained greater conformational compatibility across the reaction coordinate than the BH32.S8 single-state design, which instead tightly packed the active site around a single transition state. These findings underscore a central principle for enzyme design: productive conformational heterogeneity is an important design requirement for multistate catalysis.

Moving forward, multistate enzyme design could be combined with de novo enzyme design methods that use generative backbone design to scaffold theozymes^9,23,35^, using the newly produced protein architectures to generate conformational ensembles^36–38^ and then optimizing their active sites across the complete reaction coordinate. The same framework could be extended to enzymes undergoing large conformational changes^39,40^, provided ensembles capturing these motions are available^41,42^, allowing design to account not only for the stability of individual catalytic states but also for the pathways connecting them^43,44^. More broadly, incorporating substrate binding, intermediate rearrangement, conformational transitions and product release into multistate design objectives could address kinetic bottlenecks beyond chemical transition states. Such integration could move de novo enzyme design from its current paradigm of constructing static active sites towards the programming of dynamic protein landscapes, opening a route to efficient biocatalysts for increasingly complex new-to-nature bond-forming, bond-breaking, redox and group-transfer reactions.

## Methods

### Ensemble refinement

Ensembles were generated from crystal structures using the *phenix.ensemble_refinement* application^45^ implemented in the PHENIX software package. In ensemble refinement, electron density-restrained molecular dynamics simulations are used to sample conformational heterogeneity and generate structural ensembles consistent with the crystallographic diffraction data^20^. Prior to ensemble refinement, low-occupancy conformers were removed from the crystal structures, the remaining conformers were assigned occupancies of 1.0, and riding hydrogen atoms were added to prepare the structures for refinement. Parallel ensemble refinement simulations were performed using different combinations of the parameters p_TLS_ (0.6, 0.8, 0.9, and 1.0), τ_x_ (0.5, 1.5, and 2.0), and w_X-ray_ (2.5, 5.0, and 10.0). The parameter p_TLS_ describes the percentage of atoms included in a translation-libration-screw model used to account for global disorder, τ_x_ is the simulation time step, and w_X-ray_ is the coupled temperature-bath offset that controls the extent to which electron density contributes to the simulation force field while maintaining a target temperature of 300 K. The ensemble generated from each crystal structure with the lowest R_free_ value following ensemble refinement was selected (Supplementary Table 1).

### Theozyme placement

All calculations were performed using the Triad protein design software^24^ (Protabit, Pasadena, CA, USA). After extracting the heavy-atom coordinates from each backbone structure in the ensembles, the *addH.py* application was used to add hydrogen atoms and format atom names to the Triad convention prior to calculation. All residues in the active site, with the exception of the catalytic residues His23 and Arg124 (positions 10, 14, 19, 22, 26, 53, 64, 68, 88, 91, 92, 94, 95, 122, 123, 128, 129, 132), were mutated to Gly. Morita-Baylis-Hillman (MBH) reaction transition states (TSs) (Figure 1b)^13^ were built^18^ off of the side chain of His23 using contact geometries listed on Supplementary Table 2. Side-chain conformations of the His23 and Arg124 catalytic residues were modeled using the 2002 Dunbrack backbone-independent rotamer library with expansions of ±1 standard deviation around χ_1_ and χ ^46^. Rotamer optimization was performed using a Monte Carlo with simulated annealing search algorithm. TS pose energies were calculated using a modified version of the Phoenix energy function^24^ consisting of a Lennard-Jones 12–6 van der Waals term from the Dreiding II force field^47^ with atomic radii scaled by 0.9^48^, a direction-dependent hydrogen bond term with a well depth of 8.0 kcal mol^−1^ and an equilibrium donor−acceptor distance of 2.8 Å^48^, and an electrostatic energy term modeled using Coulomb’s law with a distance-dependent dielectric of 10. For TS1 and TS4, a standard forcefield distance cutoff of 35.0 Å was employed to capture long-range non-bonded interactions. For TS2 and TS3, the distance cutoff was reduced to 1.0 Å. This adjustment was necessary to suppress short-range steric repulsion energies (van der Waals clashes) that would otherwise occur at covalent bonding distances. A penalty energy of 1000 kcal mol^−1^ was applied when TS-side-chain interactions did not satisfy catalytic contact geometries (Supplementary Table 3). During these calculations, TS poses with energies exceeding 500 kcal mol^-1^ were discarded.

After generating the TS poses, the top placement for each transition state was selected using the following procedure. A covalent bond was introduced between TS2/TS3 and His23 in all poses generated for these transition states during theozyme placement. The energy of all four TSs was then reevaluated using the *energy.py* app in Triad and the same energy function described above to eliminate poses exhibiting steric clashes with the protein backbone. A standard forcefield distance cutoff of 35.0 Å was employed to fully capture long-range non-bonded interactions for all four TSs. Structures yielding positive energies, indicating unfavorable interactions, were discarded. For the remaining structures, the root-mean-square deviation (RMSD) between the guanidinium group of Arg124 and the imidazole group of His23 in the Triad-generated structures and their corresponding positions in the density-functional-theory-derived theozyme model (Figure 1b) was calculated. For each TS, the pose yielding the lowest RMSD was selected as the starting template for subsequent multistate and single-state enzyme design calculations, yielding four theozyme-bound structures (Supplementary Figure 1).

### Multistate enzyme design

Calculations were performed using the *multi_design.py* app in the Triad protein design software^19^. Sidechain rotamers of active-site residues (positions 10, 14, 19, 22, 26, 53, 64, 68, 88, 91, 92, 94, 95, 122, 123, 128, 129, 132), excluding the catalytic residues His23 and Arg124, were optimized on the four theozyme-bound structures, one per TS, using the sequence space listed in Supplementary Table 4. Sidechain rotamers of residues located within 4 Å of the designed positions were also optimized but their identities were fixed. The resulting sequence search space thus consisted of ∼3.4 × 10^10^ possible sequences. Rotamer optimization was carried out using the search algorithm, energy function and rotamer library described above.

In multistate design, the fitness score σ_MEnD_ to be optimized is a function of the amino acid sequence A^19^. In our calculations, the primary objective was to minimize the energy of the rate-limiting transition state (TS3) while enforcing strict stability restraints on the remaining transition states to prevent their destabilization The fitness score was calculated using an energy combination function of the following form:

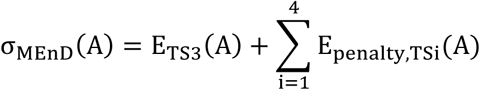

where E_TS3_(A) is the energy of sequence A bound to TS3 and E_penalty,TS i_(A) is the energy penalty applied to each TS when the energy of sequence A in the presence of this TS exceeds its corresponding reference energy.

The energy of each sequence in each state is defined as:

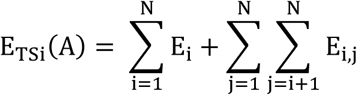

where E_i_ is the one-body energy of the amino acid at position i, corresponding to the interaction energy between its rotamer and the fixed template (i.e., the portion of the protein structure that remains unchanged), and E_i,j_ is the two-body interaction energy between pairs of rotamers at residue positions i and j.

The energy penalty was calculated according to:

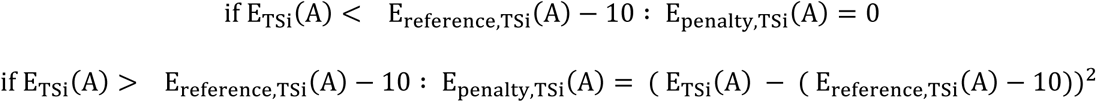

where the reference energy corresponds to the energy of the backbone template in the absence of the TS, with all designed residues, including catalytic residues, mutated to Gly. A −10 kcal mol^-1^ energy benefit was added to the reference energy to maximize stringency.

Sequence optimization was performed using a Monte Carlo search algorithm with simulated annealing, consisting of 5,000 steps with the temperature decreasing from 4,000 K to 150 K. At the end of the calculation, the 1,000 top-scoring sequences that were sampled by the Monte Carlo trajectory were subjected to a final structural cleaning and rescoring step. Cleaning involved optimizing the sidechain rotamers of all designed residues to identify the lowest-energy configuration for each sequence. The cleaned structures were then used to calculate the active-site energy for each TS, defined as the energy difference between each repacked structure containing the corresponding TS and a reference structure in which the TS and all designed residues, including catalytic residues, were mutated to Gly.

For each sequence, the difference between the active-site energies of the least-and most-stable TS was calculated to quantify the sequence’s ability to stabilize the overall reaction coordinate. Because the objective was to minimize the energy of the rate-limiting transition state, TS3, while simultaneously stabilizing the overall reaction coordinate, a cutoff active-site energy corresponding to the TS3 active-site energy of BH32.12, was applied to the TS3 active-site energies of all design outputs. Among sequences satisfying this cutoff, the 10 sequences with the smallest energy differences between transition states were selected for further analysis.

### Single-state design

Calculations were performed using the same search algorithm, energy function and rotamer library as in multistate enzyme design, with modifications to the score (σ) and the filtering criteria for active-site energy. The score corresponded to the active-site energy of the sequence in the presence of TS3, as follows:

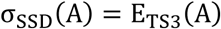

The top 10 sequences exhibiting the lowest TS3 active-site energy after cleaning were selected.

### LigandMPNN

The four theozyme-bound structures (TS1–TS4) generated as described above were merged into a single homotetramer, with each chain assigned a distinct chain ID. Sequence design was performed using LigandMPNN^26^ with the ligandmpnn_v_32_010_25.pt model weights and an 8 Å ligand cutoff distance. The redesigned positions were identical to those used in the Triad multistate design calculations (positions 10, 14, 19, 22, 26, 53, 64, 68, 88, 91, 92, 94, 95, 122, 123, 128, 129, and 132), with all 20 amino acids allowed at each position. Sampling was performed at a low temperature of 0.1 using a fixed random seed of 111 to favor high-probability amino acid assignments and ensure computational reproducibility. Fifty batches of two sequences each were generated, yielding a total of 100 sequences.

To assess structural viability, the 100 LigandMPNN-designed sequences were subjected to structure prediction using AlphaFold2^28^ with multiple sequence alignment (MSA). The predicted models were filtered using stringent confidence criteria: (i) a global mean predicted local distance difference test (pLDDT) score >95; (ii) a mean pLDDT score >95 across all redesigned positions; (iii) a pLDDT score ≥92 for every individual redesigned position; (iv) a Cα RMSD <0.8 Å relative to the BH32.12 crystal structure (PDB ID: 6Z1L); and (v) a Cα RMSD <0.8 Å for the catalytic residues His23 and Arg124, also relative to the BH32.12 crystal structure. Application of these criteria yielded 27 sequences. From these, the 10 sequences with the highest global pLDDT scores were selected for experimental testing, while requiring at least two mutations to differentiate each selected sequence from the others to favor sequence diversity.

### Protein expression and purification

Codon-optimized (*E. coli*) and His-tagged (C-terminus) genes for designed MBHases (Supplementary Table 1) cloned into the pET-29b(+) vector via *Nde*I and *Xho*I restriction sites were obtained from Twist Bioscience. Genes encoding for BH32.8 and BH32.12 encoded in plasmid pBbE8k were obtained from the laboratory of Anthony Green^13^. Enzymes were expressed in *E. coli* BL21-Gold (DE3) cells (Agilent) using lysogeny broth (LB) supplemented with 50 µg mL^−1^ kanamycin. Cultures were grown at 37 °C with shaking (250 rpm) to an optical density of 0.6–0.8 at 600 nm, after which expression was induced using 1 mM isopropyl β-D-1-thiogalactopyranoside. For BH32.8 and BH32.12, expression was instead induced using 10 mM L-arabinose. Following incubation for 18 h at 16 °C (25 °C for BH32.8 and BH32.12) with shaking at 220 rpm, cells were harvested by centrifugation (4,000 × g, 30 min), resuspended in 10 mL lysis buffer (5 mM imidazole in 100 mM potassium phosphate buffer, pH 8.0 supplemented with 1.0 mg mL^−1^ lysozyme), and lysed with an EmulsiFlex-B15 cell disruptor (Avestin). Cell debris were then removed by centrifugation (20,000 × g, 30 min), and the clarified lysates were used for protein purification.

Proteins were purified by immobilized metal affinity chromatography using Ni-NTA agarose (Qiagen) pre-equilibrated with lysis buffer in individual Econo-Pac gravity-flow columns (Bio-Rad). Bound proteins were sequentially washed with 100 mM potassium phosphate buffer (pH 8.0) containing 40 mM and 60 mM imidazole, respectively. Proteins were then eluted with the same buffer containing 250 mM imidazole. Eluted proteins were buffer-exchanged into phosphate-buffered saline (PBS) (pH 7.4) using Econo-Pac 10DG desalting prepacked gravity-flow columns (Bio-Rad). Concentrations of purified proteins were determined by measuring the absorbance at 280 nm and applying Beer-Lambert’s law using predicted extinction coefficients and molecular weights calculated from the Benchling tool (https://www.benchling.com/)

### Conversion assays

Enzyme reactions were initiated by mixing 10.5 μL of a stock solution containing 4-nitrobenzaldehyde (0.6 mM final concentration) and 2-cyclohexen-1-one (3 mM final concentration) in acetonitrile (3% v/v final concentration) with 339.5 μL of enzyme solution (20 μM final concentration) in PBS (pH 7.4) in a 2 mL HPLC vial. Reactions were carried out at 30 °C for 22 h with shaking at 800 rpm. Following incubation, reactions were quenched by the addition of 350 μL acetonitrile and shaken at 800 rpm for 2 h to precipitate the protein. Precipitated protein was removed by centrifugation (14,000 × g, 10 min), and the resulting supernatants (700 μL) were filtered through 0.2 µm microcentrifuge filter tubes (Thermo Scientific) by centrifugation (14,000 × g, 5 min) prior to HPLC analysis. All assays were performed using three independent protein batches, each measured in duplicate.

### Steady-state kinetics

Initial velocities (v_0_) were measured by varying the concentration of either 4-nitrobenzaldehyde or 2-cyclohexen-1-one while maintaining a fixed concentration of the other substrate. For measurements of v_0_ versus 4-nitrobenzaldehyde concentration, reactions contained 25 mM 2-cyclohexen-1-one and 4-nitrobenzaldehyde at 0.1–2 mM. For measurements of v_0_ versus 2-cyclohexen-1-one concentration, reactions contained 2 mM 4-nitrobenzaldehyde and 2-cyclohexen-1-one at 0.5–12.5 mM for BH32.M6 or 1–25 mM for BH32.8, BH32.12, BH32.S8 and BH32.L7. Reactions were performed with purified enzyme (90 μM BH32.S8, 30 μM BH32.8 and BH32.L7, or 10 μM BH32.12 and BH32.M6) in PBS (pH 7.4) containing 3% acetonitrile, and were incubated at 30 °C with shaking at 800 rpm. Sampling intervals were adjusted according to enzyme activity to ensure accurate determination of initial rates. Samples were quenched with 1 vol. of acetonitrile and analysed by HPLC as described below. Product concentrations were quantified using an HPLC calibration curve, and initial rates were determined by linear regression of product concentration versus time. The combined *v*_0_ versus [4-nitrobenzaldehyde] and *v*_0_ versus [2-cyclohexen-1-one] steady-state kinetic datasets were globally fitted to the random-order binding model:

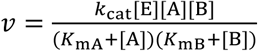

where *k*_cat_ is the catalytic constant, [E] is the total enzyme concentration, [A] and [B] are the initial 4-nitrobenzaldehyde and 2-cyclohexen-1-one concentrations, respectively, and *K*_mA_ and *K*_mB_ are the corresponding apparent Michaelis constants. Global fitting was performed by simultaneously fitting both datasets with shared *k*_cat_, *K*_mA_ and *K*_mB_.

### Kinetic isotope effects

Enzyme reactions were performed using 2-cyclohexen-1-one or 2-deuterocyclohex-2-en-1-one (10 mM), 4-nitrobenzaldehyde (2 mM) and the relevant biocatalyst (90 μM BH32.S8, 10 μM BH32.12 and BH32.M6) in PBS (pH 7.4) with 3% acetonitrile as a co-solvent. All reactions were incubated at 30 °C with shaking (800 rpm) with samples taken every 30 min for 3 h. All assays were performed in triplicate. For HPLC analysis, reactions were quenched by the addition of 1 vol. of acetonitrile and processed for HPLC analysis as described above. 2-deuterocyclohex-2-en-1-one was synthesized according to the protocol published in ^32^.

### Mechanistic inhibition

Inhibition assays were performed in PBS (pH 7.4) using a Synergy H1 microplate reader (BioTek) with 96-well UV-star microplates (Greiner Bio-One). Reactions were carried out in a final volume of 200 μL containing 4 μM enzyme and 5–100 μM inhibitor. The inhibitor was synthesized according to our previously published procedure^13^. Absorbance was recorded at 325 nm every 10 s for 30 min at 25 °C using Gen5 software (BioTek). For inhibitor assays, the time-dependent absorbance data were fitted by nonlinear regression to a one-phase association model (GraphPad Prism):

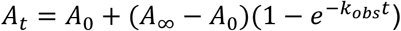

where *A_t_*is the absorbance at time *t*, *A_0_* is the initial absorbance, *A_∞_* is the plateau absorbance, and *k*_obs_ is the observed rate constant obtained from the fit.

### Kinetic solvent viscosity effects

The effects of solvent viscosity on *k*_cat_ were determined at 30 °C in PBS (pH 7.4) using sucrose as viscogen at different concentrations (0, 21, 27, 31% *w/w* for BH32.12 and BH32.M6, 0, 15, 21, 27% *w/w* for BH32.S8). Corresponding viscosities (*η*) were calculated from published viscosity data of sucrose solutions^49^. Steady-state kinetic assays were performed as described above, except that sampling intervals and total reaction times were adjusted for each enzyme variant to ensure sufficient product formation for kinetic analysis. Initial rates were determined and fitted to the Michaelis-Menten equation to calculate *k*_cat_ values. The reference value at 0% sucrose was divided by those obtained at different *η* and plotted against the relative buffer viscosity *η*_rel_ to give the corresponding slopes.

### Achiral HPLC analysis

HPLC analysis was performed on an Agilent 1100 Series Capillary LC System with a InfinityLab Poroshell 120 LC column (50 × 2.1 mm, Agilent). An isocratic mobile phase consisting of 25% acetonitrile in water was used at a flow rate of 0.5 mL min^-1^ for 5 min.

Peaks were assigned by comparison to chemically synthesized standards^13^ and the peak areas were integrated using Agilent OpenLab software.

### Enantioselectivity measurements

Analytical scale enzyme reactions were performed using 2-cyclohexen-1-one (3 mM), 4-nitrobenzaldehyde (0.6 mM) and the relevant biocatalyst (20 µM) in PBS (pH 7.4) with 3% (v/v) acetonitrile as a cosolvent at 30 °C for 24 h. Reactions were then extracted with ethyl acetate. Precipitated protein was removed by centrifugation (14,000 × g for 10 min), and the organic phase was separated and directly injected onto the SFC. Chiral analysis was performed using an SFC 1290 Infinity II system (Agilent). Enantiomers of MBH product were separated using a Daicel 80S82 CHIRALPAK IA-3 SFC column (3 mm, 50 mm and 3 µm) and an isocratic method with 20% methanol in CO_2_ at 1 mL min^−1^ for 5 min. Enantiomers of MBH product were assigned by comparison to the literature^13^. Peak areas were integrated using Agilent OpenLabs software.

### Circular dichroism (CD) and thermal denaturation assays

CD measurements were performed with a Jasco J-815 spectrometer using 300-µL aliquots of each enzyme sample at a concentration of 5 μM in 20 mM sodium phosphate buffer (pH 7.4) in a 1 mm pathlength quartz cuvette (Jasco). For structural characterization of protein folds (Supplementary Figure 3), CD spectra were acquired from 190 to 250 nm at 25°C, sampled every 1 nm at a rate of 10 nm min^-1^. Three scans were acquired and averaged for each sample. For thermal denaturation assays (Supplementary Figure 4), samples were heated at a rate of 1 °C per minute. Melting temperatures were determined by fitting the data to a two-term sigmoid function with correction for pre-and post-transition linear changes in ellipticity as a function of temperature^50^. Data were fitted to the equations below using nonlinear least squares regression in GraphPad Prism 10, where *θ*_F_ is the ellipticity when 100% folded, *θ*_U_ is the ellipticity when 100% unfolded, *c*_F_ is the linear correction for pretransition changes in ellipticity, *c*_U_ is the linear correction for post-transition changes in ellipticity, Δ*H*_U_ is the enthalpy of unfolding, *k* is the folding constant, *F* is the fraction folded, and *θ* is the ellipticity at temperature *T*.

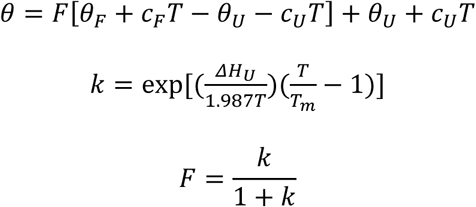

### Crystallization

For crystallographic studies, proteins obtained from His-tag affinity purification were further purified by size-exclusion chromatography on an Enrich SEC 70 column (Bio-Rad) equilibrated in 20 mM HEPES (pH 7.5) supplemented with 100 mM NaCl. Fractions containing monomeric protein were pooled and concentrated to 10–15 mg mL^−1^ using Amicon Ultra-3K centrifugal filter units (Millipore Sigma) prior to crystallization trials.

Crystals were obtained by the hanging-drop vapor diffusion method at 293 K. BH32.M6 crystals were grown in drops prepared by mixing 200 nL of protein solution (10 mg mL^−1^) with 200 nL of the mother liquor and sealing the drop inside a reservoir containing an additional 100 μL of the mother liquor solution. BH32.S8 and BH32.L7 were grown in drops prepared by mixing 1 µL of protein solution (10 mg mL^−1^) with 1 µL of the mother liquor solution. BH32.M6 crystals were grown from 0.2 M ammonium acetate, 0.1 M trisodium citrate (pH 5.6), and 30% PEG 4000; BH32.S8 crystals from 0.2 M LiCl, 0.1 M Tris (pH 9.0), and 16% PEG 6000; and BH32.L7 crystals from 0.2 M ammonium acetate, 0.1 M Na/K tartrate, 0.085 M trisodium citrate, 1.3 M ammonium sulfate, and 15% glycerol.

### X-ray data collection and processing

Crystals were mounted on polyimide loops and sealed using a MicroRT tubing kit (MiTeGen). Single-crystal X-ray diffraction data were collected on beamline 8.3.1 at the Advanced Light Source. The beamline was equipped with a Pilatus3 S 6 M detector (Dectris) and was operated at a photon energy of 11111 eV. Crystals were maintained at 280 K throughout the course of data collection. X-ray data were processed using the Xia2 software^51^, which performed indexing, integration, and scaling with DIALS^52^.

### Structure determination

Initial phase information for calculation of electron density maps was obtained by molecular replacement using the program Phaser^53^, as implemented in v1.21.1.5286 of the PHENIX suite^45^. The previously published BH32.12 structure (PDB ID: 6Z1L^13^) was used as the molecular replacement search model. BH32.M6 and BH32.S8 crystallized in the same space group *P* 1 2_1_ 1, whereas BH32.L7 crystallized in the space group *P* 2 2_1_ 2. All enzymes contain a single copy of the molecule in the crystallographic asymmetric unit. Next, we performed iterative steps of manual model rebuilding followed by refinement of atomic positions, atomic displacement parameters, and occupancies using automatic weight optimization. All model building was performed using Coot 0.9.8.92^54^ and refinement steps were performed with *phenix.refine*^55^ within the PHENIX (v1.21.1.5286) suite. Further information regarding model building and refinement are presented in Supplementary Table 8.

### Data collection and structure preparation for BH32.8 ensemble refinement

BH32.8 was purified as described previously^13^, and was prepared at a concentration of 20 mg mL^−1^ in buffer containing 50 mM HEPES and 300 mM NaCl (pH 7.5). Crystallization drops were prepared by mixing 1 µL of protein solution with 1 µL of mother liquor, followed by sealing the drop inside a reservoir containing an additional 500 µL of the mother liquor solution. The mother liquor solutions contained 0.1 M Sodium Acetate and 20% polyethylene glycol 3350. Crystals were harvested and mounted in polyimide loops and sealed in MicroRT capillaries (MiTeGen) for X-ray data collection. Single-crystal X-ray diffraction data were collected on beamline 8.3.1 at the Advanced Light Source. The beamline was equipped with a Pilatus3 S 6M detector and was operated at a photon energy of 11111 eV. Crystals were maintained at 310 K throughout the course of data collection.

X-ray data were processed with the Xia2 0.5.492 program^51^, which performed indexing, integration, and scaling with the 20180126 version of XDS and XSCALE^56^, followed by merging with Pointless as distributed in CCP4 7.0.05333^57^. Initial phase information for the calculation of electron density maps was obtained by molecular replacement using Phaser^53^, as implemented in v1.13.2998 of the PHENIX suite^55^. PDB ID 6Q7N was used as the molecular replacement search model. Several cursory rounds of model building and refinement were conducted using Coot 0.8.9.2^54^ and phenix.refine^58^, before the model was used for ensemble refinement as described above. Because the parent protein structure used to generate the ensemble is not available in the PDB, we deposited the ensemble model (PDB), and structure factors (MTZ) used to generate it, in the Zenodo database (10.5281/zenodo.22711860).

### Random-acceleration molecular dynamics (RAMD)

Conventional molecular dynamics (MD) simulations were first performed on the product-bound models of BH32.12, BH32.M6, and BH32.S8 generated using Triad^21,24^ to obtain starting structures for RAMD. Bonding parameters for the reaction product were generated using the AmberTools ANTECHAMBER^59^ module with atomic charges parametrised by RESP fitting to the HF/6-31G(d,p) electron density of a B3LYP/6-31 + G(d,p)//SCRF(water) structure optimized in Gaussian 16 Revision C.01^60^. The models were then prepared using the AmberTools^61^ TLeap module by solvating in a water box with a minimum 10 Å buffering distance around the protein and adding counter-ions. MD simulations were then carried out using Gromacs2024 with the Amber14 force field^62^, and simulations were performed using the velocity-rescaling thermostat^63^ (300 K) and Parrinello-Rahman barostat^64^ (1 bar), 10 Å van der Waals and electrostatic cut-offs, particle mesh Ewald for long-range electrostatics, LINCS bond constraints^65^, periodic boundary conditions and a 2 fs timestep. The protocol for running simulations was as follows: (i) energy minimisation; (ii) 1 ns constant volume (NVT) equilibration of the solvent with 10 kJ mol^−1^ Å^−2^ constraints on the protein; (iii) three 1 ns constant pressure (NPT) equilibration stages with 10, 1 and 0.1 kJ mol^−1^ Å^−2^ constraints; (iv) 500 ns of unconstrained production MD.

For each variant, the MD snapshot with the “most average” position of the product was selected as a starting point for RAMD simulations: the MD trajectories were aligned against the protein atoms (not including hydrogens) and the structure with the lowest RMSD for the product relative to the average was selected. A set of 20 starting structures and corresponding velocities was then generated by running a series of 5 ns MD simulations from this conformation and each was used to initiate 100 RAMD simulations for a total of 2000 per variant. The centres of mass of the product and protein were used to define the ligand-receptor pair, a force of 450 kJ mol^−1^ Å^−1^ was applied to the ligand, and the escape time was defined as the time when the centre-of-mass protein-ligand distance reached 40 Å. The resulting 2000 escape times were used to generate the distributions shown in Supplementary Figure 10. To ensure that sufficient data was used, we compared the distributions for BH32.12 and BH32.M6 obtained from a subset of 50 simulations for each of the first 10 snapshots (for a total of 500), and to test the effect of the force applied we also ran these 500 simulations with a force of 500 kJ mol^−1^ Å^−1^; in each case the same trends were observed (Supplementary Figure 12).

## Acknowledgements

R.A.C. acknowledges grants from the Natural Sciences and Engineering Research Council of Canada (RGPIN-2021-03484 and RGPAS-2021-00017) and the Canada Foundation for Innovation (26503). R.A.C., M.C.T. and A.P.G. acknowledge a joint grant from the Human Frontier Science Program (RGP0004/2022). This research was enabled in part by support provided by Compute Ontario (www.computeontario.ca) and the Digital Research Alliance of Canada (alliancecan.ca). Beamline 8.3.1 at the Advanced Light Source is operated by the University of California San Francisco with generous support from the National Institutes of Health (R01 GM124149 for technology development and P30 GM124169 for user support), and the Integrated Diffraction Analysis Technologies program of the US Department of Energy Office of Biological and Environmental Research. The Advanced Light Source (Berkeley, CA) is a national user facility operated by Lawrence Berkeley National Laboratory on behalf of the US Department of Energy under contract number DE-AC02-05CH11231, Office of Basic Energy Sciences. The contents of this publication are solely the responsibility of the authors and do not necessarily represent the official views of NIGMS or NIH. The authors thank B.J. Smith and A. Bunyat-zada for critical reading of manuscript.

## Author Contributions

Conceptualization: R.A.C and N.T.H.P. Methodology: N.T.H.P., M.C.T., A.P.G., R.A.C. Software: N.T.H.P., L.O.J. Formal analysis: N.T.H.P., R.G., A.E.C., L.O.J., J.A.W., S.H., R.A.C. Investigation: N.T.H.P., R.G., R.P.G.J., A.E.C., L.O.J., J.A.W., B.S., Z.B.-P., J.B., S.H. Resources: R.A.C., M.C.T., A.P.G. Data curation: N.T.H.P., R.G., R.A.C. Writing – original draft: R.A.C., N.T.H.P. Writing – review & editing: N.T.H.P., R.G., A.E.C., L.O.J., M.C.T., A.P.G., R.A.C. Visualization: R.A.C., N.T.H.P., R.G., A.E.C. Supervision: R.A.C. Project administration: R.A.C. Funding acquisition: R.A.C., M.C.T., A.P.G.

## Competing Interests

The authors declare no competing interests.

## Data availability

The coordinates and crystal structure factors of BH32.M6, BH32.S8 and BH32.L7 were deposited to the Protein Data Bank (PDB) under accession codes 38GE, 38GF and 38GG, respectively. The BH32.8 ensemble coordinates and corresponding structure factors (MTZ file) have been deposited in a Zenodo repository (https://zenodo.org/records/22711860). Source data are provided with this paper.

## Notes

### Competing Interest Statement

The authors have declared no competing interest.

