## Supplementary Information for "Multistate Enzyme Design Enables Efficient and Stereoselective Multistep Catalysis"

**This file includes:**

Supplementary Tables 1–9

Supplementary Figures 1–12

**Supplementary Table 1.** Ensemble refinement of MBHase structures

| | | Before ensemble refinement | | After ensemble refinement | | |
|---|---|---|---|---|---|---|
| **Enzyme** | **PDB ID** | $R_{work}$ | $R_{free}$ | $R_{work}$ | $R_{free}$ | **No. structures** |
| **BH32.8** | N.A.[a] | 0.1895 | 0.2171 | 0.1601 | 0.2047 | 40 |
| **BH32.12** | 6Z1L | 0.2230 | 0.2620 | 0.1917 | 0.2350 | 56 |
| **BH32.M6** | 38GE | 0.1768 | 0.1985 | 0.1700 | 0.1967 | 34 |
| **BH32.S8** | 38GF | 0.1834 | 0.2162 | 0.1556 | 0.2085 | 200 |
| **BH32.L7** | 38GG | 0.1723 | 0.1940 | 0.1502 | 0.1771 | 100 |

[a] This ensemble was generated from an unpublished data set of BH32.8 that we obtained at 310 K and is available at https://doi.org/10.5281/zenodo.22711860.

**Supplementary Table 2**. Geometric definitions for generation of transition-state poses relative to the His23 side chain

| TS | Residue | Type | Atom 1[a] | Atom 2[a] | Atom 3[a] | Atom 4[a] | Value[b] |
|---|---|---|---|---|---|---|---|
| **TS1** | **HID** | Distance | NE2 | **C5** | | | 1.5, 1.6, 1.7, 1.8, 1.9, 2.0, 2.1, 2.2, 2.3, 2.4, 2.5 |
| | | Angle | CE1 | NE2 | **C5** | | 117, 118, 119, 120, 121, 122, 123, 124, 125, 126, 127 |
| | | Angle | NE2 | **C5** | **C6** | | 112.8, 113.8, 114.8, 115.8, 116.8, 117.8, 118.8, 119.8, 120.8, 121.8, 122.8 |
| | | Torsion | ND1 | CE1 | NE2 | **C5** | -172.7, -171.7, -170.7, -169.7, -168.7, -167.7, -166.7, -165.7, -164.7, -163.7, -162.7 |
| | | Torsion | CE1 | NE2 | **C5** | **C6** | -8.4, -7.4, -6.4, -5.4, -4.4, -3.4, -2.4, -1.4, -0.4, 0.6, 1.6 |
| | | Torsion | NE2 | **C5** | **C6** | **C1** | -111.5, -110.5, -109.5, -108.5, -107.5, -106.5, -105.5, -104.5, -103.5, -102.5, -101.5 |
| **TS2** | **HID** | Distance | NE2 | **C5** | | | 1.3, 1.4, 1.5 |
| | | Angle | CE1 | NE2 | **C5** | | 121.9, 122.9, 123.9, 124.9, 125.6, 126.6, 127.6, 128.6, 129.6, 130.6, 131.9 |
| | | Angle | NE2 | **C5** | **C6** | | 105.4, 106.4, 107.4, 108.4, 109.4, 110.4, 111.4, 112.4, 113.4, 114.4, 115.4 |
| | | Torsion | ND1 | CE1 | NE2 | **C5** | 172.6, 173.6, 174.6, 175.6, 176.6, 177.6, 178.6, 179.6, 180.6, 181.6, 182.6 |
| | | Torsion | CE1 | NE2 | **C5** | **C6** | 34.0, 35.0, 36.0, 37.0, 38.0, 39.0, 40.0, 41.0, 42.0, 43.0, 44.0 |
| | | Torsion | NE2 | **C5** | **C6** | **C1** | -139.9, -138.9, -137.9, -136.9, -135.9, -134.9, -133.9, -132.9, -131.9, -130.9, -129.9 |
| **TS3** | **HID** | Distance | NE2 | **C5** | | | 1.3, 1.4, 1.5 |
| | | Angle | CE1 | NE2 | **C5** | | 122, 123, 124, 125, 126, 127, 128, 129, 130, 131, 132 |
| | | Angle | NE2 | **C5** | **C6** | | 106.7, 107.7, 108.7, 109.7, 110.7, 111.7, 112.7, 113.7, 114.7, 115.7, 116.7 |
| | | Torsion | ND1 | CE1 | NE2 | **C5** | 161.6, 162.6, 163.6, 164.6, 165.6, 166.6, 167.6, 168.6, 169.6, 170.6, 171.6 |
| | | Torsion | CE1 | NE2 | **C5** | **C6** | 29.9, 30.9, 31.9, 32.9, 33.9, 34.9, 35.9, 36.9, 37.9, 38.9, 39.9 |
| | | Torsion | NE2 | **C5** | **C6** | **C1** | -150.4, -149.4, -148.4, -147.4, -146.4, -145.4, -144.4, -143.4, -142.4, -141.4, -140.4 |
| **TS4** | **HID** | Distance | NE2 | **C5** | | | 1.8, 1.9, 2.0, 2.1, 2.2, 2.3, 2.4, 2.5 |
| | | Angle | CE1 | NE2 | **C5** | | 117.5, 118.5, 119.5, 120.5, 121.5, 122.5, 123.5, 124.5, 125.5, 126.5, 127.5 |
| | | Angle | NE2 | **C5** | **C6** | | 98.4, 99.4, 100.4, 101.4, 102.4, 103.4, 104.4, 105.4, 106.4, 107.4, 108.4 |
| | | Torsion | ND1 | CE1 | NE2 | **C5** | 173.3, 174.3, 175.3, 176.3, 177.3, 178.3, 179.3, 180.3, 181.3, 182.3, 183.3 |
| | | Torsion | CE1 | NE2 | **C5** | **C6** | 44.5, 45.5, 46.5, 47.5, 48.5, 49.5, 50.5, 51.5, 52.5, 53.5, 54.5 |
| | | Torsion | NE2 | **C5** | **C6** | **C1** | -100.8, -99.8, -98.8, -97.8, -96.8, -95.8, -94.8, -93.8, -92.8, -91.8, -90.8 |

[a] Atoms in bold are from the transition state. All other atoms are from the catalytic residue.
[b] Distance measurements given in Å, all others are in degrees.

**Supplementary Table 3**. Geometric constraints used to define catalytic contacts

| TS | Residue | Type | Atom 1[a] | Atom 2[a] | Atom 3[a] | Atom 4[a] | Min[b] | Max[b] |
|---|---|---|---|---|---|---|---|---|
| **TS1** | **His23** | Distance | NE2 | **C5** | | | 1.5 | 2.5 |
| | | Angle | CE1 | NE2 | **C5** | | 117 | 127 |
| | | Angle | NE2 | **C5** | **C6** | | 112.8 | 122.8 |
| | | Torsion | ND1 | CE1 | NE2 | **C5** | -172.7 | -162.7 |
| | | Torsion | CE1 | NE2 | **C5** | **C6** | -8.4 | 1.6 |
| | | Torsion | NE2 | **C5** | **C6** | **C1** | -111.5 | -101.5 |
| | **Arg124** | Distance | NH1 | **O1** | | | 2.6 | 3.0 |
| | | Distance | NH2 | **O1** | | | 2.6 | 3.0 |
| | | Distance | NH2 | **O7** | | | 2.8 | 3.2 |
| | | Angle | **O1** | NH1 | NH2 | | 55.5 | 75.1 |
| | | Angle | **O7** | **O1** | NH1 | | 77.2 | 97.2 |
| | | Torsion | **O1** | NH1 | NH2 | CZ | 161.1 | 181.1 |
| | | Torsion | **O7** | **O1** | NH1 | NH2 | 4.4 | 24.4 |
| | | Torsion | **C1** | **O7** | **O1** | NH1 | -179.1 | -159.7 |
| **TS2** | **His23** | Distance | NE2 | **C5** | | | 1.3 | 1.5 |
| | | Angle | CE1 | NE2 | **C5** | | 121.9 | 131.9 |
| | | Angle | NE2 | **C5** | **C6** | | 105.4 | 115.4 |
| | | Torsion | ND1 | CE1 | NE2 | **C5** | 172.6 | 182.6 |
| | | Torsion | CE1 | NE2 | **C5** | **C6** | 34 | 44 |
| | | Torsion | NE2 | **C5** | **C6** | **C1** | -139.9 | -129.9 |
| | **Arg124** | Distance | NH1 | **O1** | | | 2.5 | 2.9 |
| | | Distance | NH2 | **O1** | | | 3.2 | 3.6 |
| | | Distance | NH2 | **O7** | | | 2.5 | 2.9 |
| | | Angle | **O1** | NH1 | NH2 | | 74 | 94 |
| | | Angle | **O7** | **O1** | NH1 | | 64.9 | 84.9 |
| | | Torsion | **O1** | NH1 | NH2 | CZ | 124 | 144 |
| | | Torsion | **O7** | **O1** | NH1 | NH2 | 32.5 | 52.5 |
| | | Torsion | **C1** | **O7** | **O1** | NH1 | 158.1 | 178.1 |
| **TS3** | **His23** | Distance | NE2 | **C5** | | | 1.3 | 1.5 |
| | | Angle | CE1 | NE2 | **C5** | | 122.0 | 132.0 |
| | | Angle | NE2 | **C5** | **C6** | | 106.7 | 116.7 |
| | | Torsion | ND1 | CE1 | NE2 | **C5** | 116.6 | 171.6 |
| | | Torsion | CE1 | NE2 | **C5** | **C6** | 29.9 | 39.9 |
| | | Torsion | NE2 | **C5** | **C6** | **C1** | -150.4 | -140.4 |
| | **Arg124** | Distance | NH1 | **O1** | | | 2.5 | 2.9 |
| | | Distance | NH2 | **O1** | | | 2.6 | 3.0 |
| | | Distance | NH2 | **O7** | | | 3.0 | 3.4 |
| | | Angle | **O1** | NH1 | NH2 | | 68.1 | 88.1 |
| | | Angle | **O7** | **O1** | NH1 | | 73.2 | 93.2 |
| | | Torsion | **O1** | NH1 | NH2 | CZ | 130.6 | 150.6 |
| | | Torsion | **O7** | **O1** | NH1 | NH2 | 33.5 | 53.5 |
| | | Torsion | **C1** | **O7** | **O1** | NH1 | 123.4 | 153.4 |
| **TS4** | **His23** | Distance | NE2 | **C5** | | | 1.8 | 2.5 |
| | | Angle | CE1 | NE2 | **C5** | | 117.5 | 127.5 |
| | | Angle | NE2 | **C5** | **C6** | | 98.4 | 108.4 |
| | | Torsion | ND1 | CE1 | NE2 | **C5** | 173.3 | 183.3 |
| | | Torsion | CE1 | NE2 | **C5** | **C6** | 44.5 | 54.5 |
| | | Torsion | NE2 | **C5** | **C6** | **C1** | -100.8 | -90.8 |
| | **Arg124** | Distance | NH1 | **O1** | | | 2.6 | 3.0 |
| | | Distance | NH2 | **O1** | | | 2.6 | 3.0 |
| | | Angle | **O1** | NH1 | NH2 | | 54.9 | 74.9 |
| | | Angle | **O7** | **O1** | NH1 | | 121.4 | 141.4 |
| | | Torsion | **O1** | NH1 | NH2 | CZ | 163.5 | 183.5 |
| | | Torsion | **O7** | **O1** | NH1 | NH2 | 20.6 | 40.6 |
| | | Torsion | **C1** | **O7** | **O1** | NH1 | -180.1 | -160.1 |

[a] Atoms in bold are from the transition state. All other atoms are from the catalytic residue.
[b] Distance measurements given in Å, all others are in degrees.

**Supplementary Table 4.** Amino-acid positions optimized during multistate and single-state active-site repacking

| Designed positions | Allowed amino acids [a] | BH32.12 amino acids |
|---|---|---|
| **10** | **TRP**, PHE, TYR | TRP |
| **14** | ILE, LEU, MET, ALA, VAL, TRP, PHE, TYR, **THR** | ILE |
| **19** | **THR** | ALA |
| **22** | ILE, LEU, ALA, **VAL**, TRP, PHE, TYR | VAL |
| **26** | **ILE**, LEU, ALA, VAL, TRP, PHE, TYR | ILE |
| **53** | **ILE**, LEU, ALA, VAL, TRP, PHE, TYR | PHE |
| **64** | ILE, LEU, **ALA**, VAL, TRP, PHE, TYR | LEU |
| **68** | ILE, LEU, ALA, VAL, TRP, **PHE**, TYR | LEU |
| **88** | ILE, LEU, MET, **ALA**, VAL, TRP, PHE, TYR | TRP |
| **91** | **SER** | SER |
| **92** | ILE, LEU, **MET**, ALA, VAL, TRP, PHE, TYR | LEU |
| **94** | **MET** | MET |
| **95** | ILE, LEU, MET, **ALA**, VAL, TRP, PHE, TYR | ALA |
| **122** | ILE, LEU, ALA, VAL, TRP, **PHE**, TYR | LEU |
| **123** | **ASN** | ASN |
| **128** | ILE, **LEU**, ALA, VAL, TRP, PHE, TYR | PRO |
| **129** | **SER** | SER |
| **132** | TRP, **PHE**, TYR | PHE |

[a] Mutations in the BH32.M6 sequence are highlighted in bold, and mutations in the BH32.S8 sequence are marked with underlining.

**Supplementary Table 5.** Amino-acid sequences of BH32 family MBHases

| Enzyme | MW [a] (kDa) | Sequence |
|---|---|---|
| **BH32.8** | 27.6 | MIRAVFFDSLGTLISVEGAYKVHLKIMEEVLGDYPLNPKTLLDEYEKLAREAFSNYAGKPYRPLRDILEEVM<br>RKLAEKYGFKYPENLWEISLRMAQRYGELYPEVVEVLKSLKGKYHVGVILNRDTEPATAFLDALGIKDLFD<br>SITTSEEAGFFKPHPRIFELALKKAGVKGEKAVYVGDNPVKDAGGSKNLGMTSILLDRKGEKREFWDKADF<br>IVSDLREVIKIVDELNGQGSLEHHHHHH* |
| **BH32.12** | 28.6 | MIRAVFFDSWGTLISVEGAYKVHFKIMEEVLGDYPLNPKTLLDEYEKLAREAFSNNAGKPYRPLRDILEEVM<br>RKLAEKYGFKYPENLWEISLRMAQRYGELYPEVVEVLKSLKGKYHVGVILNRDTEPSTAFLDALGIKDLFD<br>SITTSEEAGFSKPHPRIFELALKKAGVKGEKAVCVGPNPVKDAGGSKNLGMTSILLDRKGEKREFWDKAD<br>FIVSDLREVIKIVDELNGQGSLEWSHPQFEKHHHHHH* |
| **BH32.M1** | 28.5 | MIRAVFFDSWGTLTSVEGTYKVHFKIMEEVLGDYPLNPKTLLDEYEKLAREAVSNNAGKPYRPWRDIIEEV<br>MRKLAEKYGFKYPENLAEISLRMAQRYGELYPEVVEVLKSLKGKYHVGVIFNRDTELSTAFLDALGIKDLFD<br>SITTSEEAGFSKPHPRIFELALKKAGVKGEKAVCVGPNPVKDAGGSKNLGMTSILLDRKGEKREFWDKAD<br>FIVSDLREVIKIVDELNGQGSLEWSHPQFEKHHHHHH* |
| **BH32.M2** | 28.5 | MIRAVFFDSWGTLMSVEGTYKVHFKIMEEVLGDYPLNPKTLLDEYEKLAREAISNNAGKPYRPWRDIIEEV<br>MRKLAEKYGFKYPENLAEISARMAQRYGELYPEVVEVLKSLKGKYHVGVIFNRDTELSTAFLDALGIKDLFD<br>SITTSEEAGFSKPHPRIFELALKKAGVKGEKAVCVGPNPVKDAGGSKNLGMTSILLDRKGEKREFWDKAD<br>FIVSDLREVIKIVDELNGQGSLEWSHPQFEKHHHHHH* |
| **BH32.M3** | 28.5 | MIRAVFFDSWGTLTSVEGTYKVHFKIMEEVLGDYPLNPKTLLDEYEKLAREAISNNAGKPYRPARDIYEEV<br>MRKLAEKYGFKYPENLAEISMRMAQRYGELYPEVVEVLKSLKGKYHVGVIFNRDTELSTAFLDALGIKDLF<br>DSITTSEEAGFSKPHPRIFELALKKAGVKGEKAVCVGPNPVKDAGGSKNLGMTSILLDRKGEKREFWDKA<br>DFIVSDLREVIKIVDELNGQGSLEWSHPQFEKHHHHHH* |
| **BH32.M4** | 28.4 | MIRAVFFDSWGTLVSVEGTYKVHFKIMEEVLGDYPLNPKTLLDEYEKLAREAISNNAGKPYRPARDIFEEV<br>MRKLAEKYGFKYPENLAEISMRMAQRYGELYPEVVEVLKSLKGKYHVGVIFNRDTEASTAFLDALGIKDLF<br>DSITTSEEAGFSKPHPRIFELALKKAGVKGEKAVCVGPNPVKDAGGSKNLGMTSILLDRKGEKREFWDKA<br>DFIVSDLREVIKIVDELNGQGSLEWSHPQFEKHHHHHH* |
| **BH32.M5** | 28.5 | MIRAVFFDSWGTLTSVEGTYKVHFKIMEEVLGDYPLNPKTLLDEYEKLAREAISNNAGKPYRPWRDILEEV<br>MRKLAEKYGFKYPENLAEISIRMAQRYGELYPEVVEVLKSLKGKYHVGVIINRDTELSTAFLDALGIKDLFDS<br>ITTSEEAGFSKPHPRIFELALKKAGVKGEKAVCVGPNPVKDAGGSKNLGMTSILLDRKGEKREFWDKADFI<br>VSDLREVIKIVDELNGQGSLEWSHPQFEKHHHHHH* |
| **BH32.M6** | 28.5 | MIRAVFFDSWGTLTSVEGTYKVHFKIMEEVLGDYPLNPKTLLDEYEKLAREAISNNAGKPYRPARDIFEEV<br>MRKLAEKYGFKYPENLAEISMRMAQRYGELYPEVVEVLKSLKGKYHVGVIFNRDTELSTAFLDALGIKDLF<br>DSITTSEEAGFSKPHPRIFELALKKAGVKGEKAVCVGPNPVKDAGGSKNLGMTSILLDRKGEKREFWDKA<br>DFIVSDLREVIKIVDELNGQGSLEWSHPQFEKHHHHHH* |
| **BH32.M7** | 28.6 | MIRAVFFDSWGTLMSVEGTYKVHFKLMEEVLGDYPLNPKTLLDEYEKLAREAISNNAGKPYRPWRDIIEEV<br>MRKLAEKYGFKYPENLAEISMRMAQRYGELYPEVVEVLKSLKGKYHVGVIFNRDTELSTAFLDALGIKDLF<br>DSITTSEEAGFSKPHPRIFELALKKAGVKGEKAVCVGPNPVKDAGGSKNLGMTSILLDRKGEKREFWDKA<br>DFIVSDLREVIKIVDELNGQGSLEWSHPQFEKHHHHHH* |
| **BH32.M8** | 28.5 | MIRAVFFDSWGTLASVEGTYKVHFKIMEEVLGDYPLNPKTLLDEYEKLAREAISNNAGKPYRPARDIYEEV<br>MRKLAEKYGFKYPENLAEISMRMAQRYGELYPEVVEVLKSLKGKYHVGVIFNRDTELSTAFLDALGIKDLF<br>DSITTSEEAGFSKPHPRIFELALKKAGVKGEKAVCVGPNPVKDAGGSKNLGMTSILLDRKGEKREFWDKA<br>DFIVSDLREVIKIVDELNGQGSLEWSHPQFEKHHHHHH* |
| **BH32.M9** | 28.7 | MIRAVFFDSWGTLTSVEGTYKVHFKIMEEVLGDYPLNPKTLLDEYEKLAREAISNNAGKPYRPWRDIIEEV<br>MRKLAEKYGFKYPENLMEISMRMAQRYGELYPEVVEVLKSLKGKYHVGVIFNRDTEWSTAFLDALGIKDLF<br>DSITTSEEAGFSKPHPRIFELALKKAGVKGEKAVCVGPNPVKDAGGSKNLGMTSILLDRKGEKREFWDKA<br>DFIVSDLREVIKIVDELNGQGSLEWSHPQFEKHHHHHH* |
| **BH32.M10** | 28.5 | MIRAVFFDSWGTLTSVEGTYKVHFKIMEEVLGDYPLNPKTLLDEYEKLAREAISNNAGKPYRPWRDIIEEV<br>MRKLAEKYGFKYPENLAEISMRMAQRYGELYPEVVEVLKSLKGKYHVGVIFNRDTEASTAFLDALGIKDLF<br>DSITTSEEAGFSKPHPRIFELALKKAGVKGEKAVCVGPNPVKDAGGSKNLGMTSILLDRKGEKREFWDKA<br>DFIVSDLREVIKIVDELNGQGSLEWSHPQFEKHHHHHH* |
| **BH32.S1** | 28.6 | MIRAVFFDSWGTLASVEGTYKVHFKFMEEVLGDYPLNPKTLLDEYEKLAREAISNNAGKPYRPWRDIIEEV<br>MRKLAEKYGFKYPENLAEISMRMMQRYGELYPEVVEVLKSLKGKYHVGVIFNRDTELSTAFLDALGIKDLF<br>DSITTSEEAGFSKPHPRIFELALKKAGVKGEKAVCVGPNPVKDAGGSKNLGMTSILLDRKGEKREFWDKA<br>DFIVSDLREVIKIVDELNGQGSLEWSHPQFEKHHHHHH* |
| **BH32.S2** | 28.7 | MIRAVFFDSWGTLTSVEGTYKVHFKFMEEVLGDYPLNPKTLLDEYEKLAREAFSNNAGKPYRPWRDIIEEV<br>MRKLAEKYGFKYPENLAEISMRMMQRYGELYPEVVEVLKSLKGKYHVGVIFNRDTELSTAFLDALGIKDLF<br>DSITTSEEAGFSKPHPRIFELALKKAGVKGEKAVCVGPNPVKDAGGSKNLGMTSILLDRKGEKREFWDKA<br>DFIVSDLREVIKIVDELNGQGSLEWSHPQFEKHHHHHH* |

| | | |
|---|---|---|
| **BH32.S3** | 28.7 | MIRAVFFDSWGTLTSVEGTYKVHFKFMEEVLGDYPLNPKTLLDEYEKLAREAISNNAGKPYRPWRDIIEEV<br>MRKLAEKYGFKYPENLAEISMRMMQRYGELYPEVVEVLKSLKGKYHVGVIFNRDTELSTAYLDALGIKDLF<br>DSITTSEEAGFSKPHPRIFELALKKAGVKGEKAVCVGPNPVKDAGGSKNLGMTSILLDRKGEKREFWDKA<br>DFIVSDLREVIKIVDELNGQGSLEWSHPQFEKHHHHHH* |
| **BH32.S4** | 28.5 | MIRAVFFDSWGTLTSVEGTYKVHFKFMEEVLGDYPLNPKTLLDEYEKLAREAISNNAGKPYRPARDILEEV<br>MRKLAEKYGFKYPENLAEISMRMMQRYGELYPEVVEVLKSLKGKYHVGVIFNRDTELSTAFLDALGIKDLF<br>DSITTSEEAGFSKPHPRIFELALKKAGVKGEKAVCVGPNPVKDAGGSKNLGMTSILLDRKGEKREFWDKA<br>DFIVSDLREVIKIVDELNGQGSLEWSHPQFEKHHHHHH* |
| **BH32.S5** | 28.6 | MIRAVFFDSWGTLTSVEGTYKVHFKFMEEVLGDYPLNPKTLLDEYEKLAREAISNNAGKPYRPWRDIIEEV<br>MRKLAEKYGFKYPENLAEISMRMIQRYGELYPEVVEVLKSLKGKYHVGVIFNRDTELSTAFLDALGIKDLFD<br>SITTSEEAGFSKPHPRIFELALKKAGVKGEKAVCVGPNPVKDAGGSKNLGMTSILLDRKGEKREFWDKAD<br>FIVSDLREVIKIVDELNGQGSLEWSHPQFEKHHHHHH* |
| **BH32.S6** | 28.7 | MIRAVFFDSWGTLTSVEGTYKVHFKFMEEVLGDYPLNPKTLLDEYEKLAREALSNNAGKPYRPWRDIIEEV<br>MRKLAEKYGFKYPENLAEISMRMMQRYGELYPEVVEVLKSLKGKYHVGVIFNRDTELSTAFLDALGIKDLF<br>DSITTSEEAGFSKPHPRIFELALKKAGVKGEKAVCVGPNPVKDAGGSKNLGMTSILLDRKGEKREFWDKA<br>DFIVSDLREVIKIVDELNGQGSLEWSHPQFEKHHHHHH* |
| **BH32.S7** | 28.7 | MIRAVFFDSWGTLMSVEGTYKVHFKFMEEVLGDYPLNPKTLLDEYEKLAREAISNNAGKPYRPWRDIIEEV<br>MRKLAEKYGFKYPENLAEISMRMIQRYGELYPEVVEVLKSLKGKYHVGVIFNRDTELSTAYLDALGIKDLFD<br>SITTSEEAGFSKPHPRIFELALKKAGVKGEKAVCVGPNPVKDAGGSKNLGMTSILLDRKGEKREFWDKAD<br>FIVSDLREVIKIVDELNGQGSLEWSHPQFEKHHHHHH* |
| **BH32.S8** | 28.6 | MIRAVFFDSWGTLASVEGTYKVHFKFMEEVLGDYPLNPKTLLDEYEKLAREAISNNAGKPYRPWRDIIEEV<br>MRKLAEKYGFKYPENLAEISMRMIQRYGELYPEVVEVLKSLKGKYHVGVIFNRDTELSTAYLDALGIKDLFD<br>SITTSEEAGFSKPHPRIFELALKKAGVKGEKAVCVGPNPVKDAGGSKNLGMTSILLDRKGEKREFWDKAD<br>FIVSDLREVIKIVDELNGQGSLEWSHPQFEKHHHHHH* |
| **BH32.S9** | 28.6 | MIRAVFFDSFGTLTSVEGTYKVHFKFMEEVLGDYPLNPKTLLDEYEKLAREAISNNAGKPYRPWRDIIEEV<br>MRKLAEKYGFKYPENLAEISMRMMQRYGELYPEVVEVLKSLKGKYHVGVIFNRDTEASTAFLDALGIKDLF<br>DSITTSEEAGFSKPHPRIFELALKKAGVKGEKAVCVGPNPVKDAGGSKNLGMTSILLDRKGEKREFWDKA<br>DFIVSDLREVIKIVDELNGQGSLEWSHPQFEKHHHHHH* |
| **BH32.S10** | 28.6 | MIRAVFFDSWGTLTSVEGTYKVHFKFMEEVLGDYPLNPKTLLDEYEKLAREAVSNNAGKPYRPWRDIIEEV<br>MRKLAEKYGFKYPENLAEISMRMMQRYGELYPEVVEVLKSLKGKYHVGVIFNRDTELSTAFLDALGIKDLF<br>DSITTSEEAGFSKPHPRIFELALKKAGVKGEKAVCVGPNPVKDAGGSKNLGMTSILLDRKGEKREFWDKA<br>DFIVSDLREVIKIVDELNGQGSLEWSHPQFEKHHHHHH* |
| **BH32.L1** | 27.1 | MIRAVFFDSLGTLLSVEGAYKVHFKVMEEVLGDYPLNPKTLLDEYEKLAREARSNNAGKPYRPDRDIREEV<br>MRKLAEKYGFKYPENLMEILQRMAQRYGELYPEVVEVLKSLKGKYHVGVIVDRDTEMTTAGLDALGIKDLF<br>DSITTSEEAGFSKPHPRIFELALKKAGVKGEKAVCVGPNPVKDAGGSKNLGMTSILLDRKGEKREFWDKA<br>DFIVSDLREVIKIVDELNGLEHHHHHH* |
| **BH32.L2** | 27.1 | MIRAVFFDSLGTLLSVEGAYKVHFKVMEEVLGDYPLNPKTLLDEYEKLAREARSNNAGKPYRPTRDIDEEV<br>MRKLAEKYGFKYPENLEEILRRMAQRYGELYPEVVEVLKSLKGKYHVGVIVDRDTEMTTAGLDALGIKDLF<br>DSITTSEEAGFSKPHPRIFELALKKAGVKGEKAVCVGPNPVKDAGGSKNLGMTSILLDRKGEKREFWDKA<br>DFIVSDLREVIKIVDELNGLEHHHHHH* |
| **BH32.L3** | 27.2 | MIRAVFFDSLGTLLSVEGAYKVHFKVMEEVLGDYPLNPKTLLDEYEKLAREAYSNNAGKPYRPTRDIDEEV<br>MRKLAEKYGFKYPENLWEIKQRMAQRYGELYPEVVEVLKSLKGKYHVGVIVDRDTEETTAGLDALGIKDLF<br>DSITTSEEAGFSKPHPRIFELALKKAGVKGEKAVCVGPNPVKDAGGSKNLGMTSILLDRKGEKREFWDKA<br>DFIVSDLREVIKIVDELNGLEHHHHHH* |
| **BH32.L4** | 27.1 | MIRAVFFDSLGTLLSVEGAYKVHFKVMEEVLGDYPLNPKTLLDEYEKLAREAYSNNAGKPYRPTRDIDEEV<br>MRKLAEKYGFKYPENLEEILQRMAQRYGELYPEVVEVLKSLKGKYHVGVIVDRDTEMTTAGLDALGIKDLF<br>DSITTSEEAGFSKPHPRIFELALKKAGVKGEKAVCVGPNPVKDAGGSKNLGMTSILLDRKGEKREFWDKA<br>DFIVSDLREVIKIVDELNGLEHHHHHH* |
| **BH32.L5** | 27.1 | MIRAVFFDSLGTLLSVEGAYKVHFKVMEEVLGDYPLNPKTLLDEYEKLAREAYSNNAGKPYRPTRDIDEEV<br>MRKLAEKYGFKYPENLMEILRRMAQRYGELYPEVVEVLKSLKGKYHVGVIVDRDTEDTTAGLDALGIKDLF<br>DSITTSEEAGFSKPHPRIFELALKKAGVKGEKAVCVGPNPVKDAGGSKNLGMTSILLDRKGEKREFWDKA<br>DFIVSDLREVIKIVDELNGLEHHHHHH* |
| **BH32.L6** | 27.2 | MIRAVFFDSLGTLLSVEGAYKVHFKVMEEVLGDYPLNPKTLLDEYEKLAREAYSNNAGKPYRPTRDIHEEV<br>MRKLAEKYGFKYPENLWEILQRMAQRYGELYPEVVEVLKSLKGKYHVGVIVDRDTEETTAGLDALGIKDLF<br>DSITTSEEAGFSKPHPRIFELALKKAGVKGEKAVCVGPNPVKDAGGSKNLGMTSILLDRKGEKREFWDKA<br>DFIVSDLREVIKIVDELNGLEHHHHHH* |
| **BH32.L7** | 27.2 | MIRAVFFDSLGTLLSVEGAYKVHFKVMEEVLGDYPLNPKTLLDEYEKLAREAYSNNAGKPYRPTRDIHEEV<br>MRKLAEKYGFKYPENLWEILQRMVQRYGELYPEVVEVLKSLKGKYHVGVIVDRDTEMTTAALDALGIKDLF<br>DSITTSEEAGFSKPHPRIFELALKKAGVKGEKAVCVGPNPVKDAGGSKNLGMTSILLDRKGEKREFWDKA<br>DFIVSDLREVIKIVDELNGLEHHHHHH* |

| | | |
|---|---|---|
| **BH32.L8** | 27.1 | MIRAVFFDSLGTLLSVEGAYKVHFKVMEEVLGDYPLNPKTLLDEYEKLAREAYSNNAGKPYRPTRDIREEV<br>MRKLAEKYGFKYPENLEEILLRMAQRYGELYPEVVEVLKSLKGKYHVGVILDRDTEMSTAGLDALGIKDLF<br>DSITTSEEAGFSKPHPRIFELALKKAGVKGEKAVCVGPNPVKDAGGSKNLGMTSILLDRKGEKREFWDKA<br>DFIVSDLREVIKIVDELNGLEHHHHHH* |
| **BH32.L9** | 27.1 | MIRAVFFDSLGTLLSVEGAYKIHFKVMEEVLGDYPLNPKTLLDEYEKLAREAYSNNAGKPYRPTRDIREEV<br>MRKLAEKYGFKYPENLIEILQRMAQRYGELYPEVVEVLKSLKGKYHVGVIVDRDTELTTAGLDALGIKDLFD<br>SITTSEEAGFSKPHPRIFELALKKAGVKGEKAVCVGPNPVKDAGGSKNLGMTSILLDRKGEKREFWDKAD<br>FIVSDLREVIKIVDELNGLEHHHHHH* |
| **BH32.L10** | 27.1 | MIRAVFFDSLGTLLSVEGSYKVHFKVMEEVLGDYPLNPKTLLDEYEKLAREAYSNNAGKPYRPTRDIDEEV<br>MRKLAEKYGFKYPENLMEILQRMAQRYGELYPEVVEVLKSLKGKYHVGVIVDRDTEMTTAGLDALGIKDLF<br>DSITTSEEAGFSKPHPRIFELALKKAGVKGEKAVCVGPNPVKDAGGSKNLGMTSILLDRKGEKREFWDKA<br>DFIVSDLREVIKIVDELNGLEHHHHHH* |

[a] Molecular weight

**Supplementary Table 6.** Characterization of enzyme variants

| Enzyme | Yield (mg $L^{-1}$) [a] | $T_m$ (°C) [b] |
|---|---|---|
| **BH32.8** | 80 ± 30 | 81.5 ± 0.2 |
| **BH32.12** | 90 ± 20 | 68.9 ± 0.2 |
| **BH32.M1** | 60 ± 20 | 69.8 ± 0.3 |
| **BH32.M2** | 50 ± 10 | 69.6 ± 0.1 |
| **BH32.M3** | 50 ± 8 | 65.9 ± 0.1 |
| **BH32.M4** | 26 ± 4 | 66.4 ± 0.1 |
| **BH32.M5** | 28 ± 2 | 70.3 ± 0.3 |
| **BH32.M6** | 57 ± 5 | 66.8 ± 0.1 |
| **BH32.M7** | 40 ± 4 | 69.2 ± 0.2 |
| **BH32.M8** | 40 ± 8 | 68.2 ± 0.1 |
| **BH32.M9** | 27 ± 1 | 69.6 ± 0.2 |
| **BH32.M10** | 30 ± 3 | 64.24 ± 0.08 |
| **BH32.S1** | 39 ± 8 | 67.7 ± 0.1 |
| **BH32.S2** | 18 ± 5 | 64.2 ± 0.1 |
| **BH32.S3** | 39 ± 4 | 64.4 ± 0.2 |
| **BH32.S4** | 28 ± 7 | 65.5 ± 0.3 |
| **BH32.S5** | 40 ± 10 | 63.7 ± 0.2 |
| **BH32.S6** | 30 ± 13 | 63.2 ± 0.1 |
| **BH32.S7** | 45 ± 5 | 68.46 ± 0.09 |
| **BH32.S8** | 30 ± 3 | 65.73 ± 0.09 |
| **BH32.S9** | 50 ± 10 | 69.0 ± 0.1 |
| **BH32.S10** | 36 ± 3 | 63.14 ± 0.08 |
| **BH32.L1** | 40 ± 10 | 69.6 ± 0.1 |
| **BH32.L2** | 9 ± 3 | 78.1 ± 0.8 |
| **BH32.L3** | 50 ± 20 | 71.0 ± 0.2 |
| **BH32.L4** | 40 ± 30 | 65.4 ± 0.8 |
| **BH32.L5** | 8 ± 1 | 55 ± 3 |
| **BH32.L6** | 15 ± 5 | 61 ± 2 |
| **BH32.L7** | 40 ± 9 | 75.9 ± 0.8 |
| **BH32.L8** | 20 ± 8 | 64 ± 2 |
| **BH32.L9** | 50 ± 30 | 68.0 ± 0.1 |
| **BH32.L10** | 50 ± 30 | 59.2 ± 0.9 |

[a] Protein expression and purification yield (mean ± s.d., n = 2–7 individual protein batches).
[b] Melting temperatures determined from a single protein batch (n = 1), with errors representing the standard errors obtained from the curve fitting.

**Supplementary Table 7.** Crystallization conditions

| Enzyme[a] | Protein (mg mL$^{-1}$) | Buffer |
|---|---|---|
| **BH32.M6** | 10 | 0.2 M ammonium acetate<br>0.1 M tri-sodium citrate pH 5.6<br>30% PEG-4000 |
| **BH32.S8** | 10 | 0.2 M LiCl<br>0.1M Tris pH 9.0<br>16% PEG-6000 |
| **BH32.L7** | 10 | 0.17 M Na/K Tartrate<br>0.085 M tri-sodium citrate<br>1.3M ammonium sulfate<br>15% glycerol |

[a] All proteins were crystallized at 20 °C.

**Supplementary Table 8.** Crystallography data and refinement statistics

| | **BH32.M6** | **BH32.S8** | **BH32.L7** |
|---|---|---|---|
| PDB ID | 38GE | 38GF | 38GG |
| **Data collection** [a] | | | |
| Temperature (K) | 280 | 280 | 280 |
| Resolution (Å) | 69.52–2.17 | 69.56–1.90 | 64.50–1.80 |
| Space group | $P$ 1 $2_1$ 1 | $P$ 1 $2_1$ 1 | $P$ 2 $2_1$ 2 |
| *Cell params.* | | | |
| a b c (Å) | 35.196<br>69.626<br>56.436 | 35.227<br>69.556<br>56.293 | 34.477<br>70.538<br>159.736 |
| α β γ (°) | 90<br>103.323<br>90 | 90<br>103.528<br>90 | 90<br>90<br>90 |
| Chains per asymm. unit | 1 | 1 | 1 |
| $R_{pim}$ | 0.059 (0.565) | 0.037 (0.446) | 0.048 (0.821) |
| $CC_{1/2}$ | 0.996 (0.440) | 0.998 (0.704) | 0.992 (0.585) |
| I/σI | 9.6 (1.2) | 11.3 (0.8) | 8.6 (0.8) |
| Completeness (%) | 98.7 (98.5) | 98.1 (95.7) | 96.4 (98.1) |
| Multiplicity | 3.3 (3.2) | 3.3 (3.4) | 3.3 (3.0) |
| Wilson B-factor (Å$^2$) | 35.480 | 31.050 | 25.590 |
| # unique reflections | 13873 (700) | 20523 (992) | 35839 (1780) |
| **Refinement** | | | |
| R work/free | 0.1768/0.1985 | 0.1834/0.2162 | 0.1711/0.1928 |
| *No. atoms* | | | |
| Protein | 1881 | 1920 | 2068 |
| Ligand | 1 | 1 | 5 |
| Water | 29 | 48 | 68 |
| *Averaged B-factors (Å$^2$)* | | | |
| Protein | 49.16 | 46.01 | 37.37 |
| Ligands | 36.74 | 27.75 | 30.88 |
| Water | 47.36 | 42.75 | 43.92 |
| *RMSD* | | | |
| bond lengths (Å) | 0.003 | 0.004 | 0.007 |
| bond angles (°) | 0.570 | 0.640 | 0.790 |
| *Molprobity statistics* | | | |
| Ramachand. outliers (%) | 0.00 | 0.00 | 0.00 |
| Ramachand. allowed (%) | 2.62 | 3.06 | 1.27 |
| Ramachan. favored (%) | 97.38 | 96.94 | 98.73 |
| Rotamer outliers (%) | 0.00 | 0.00 | 0.00 |
| MolProbity clashscore | 1.85 | 2.86 | 3.38 |

[a] Highest resolution shell is shown in parentheses.

**Supplementary Table 9.** Inhibition of various MBHases

| Enzyme | $k_{inact}$ ($s^{-1}$) | $K_I$ (μM) | $k_{inact}/K_I$ ($M^{-1}$ $s^{-1}$) |
|---|---|---|---|
| BH32.M6 | 0.045 ± 0.005 | 90 ± 20 | 500 |
| BH32.S8 | 0.011 ± 0.002 | 180 ± 40 | 60 |

Kinetic assays were carried out at 25 °C in phosphate buffer (pH 7.4) containing 3% acetonitrile (v/v). For BH32.M6, two independent protein batches were analyzed with four and six technical replicates, respectively; for BH32.S8, two independent protein batches were analyzed with six technical replicates each. Errors on the fit are provided.

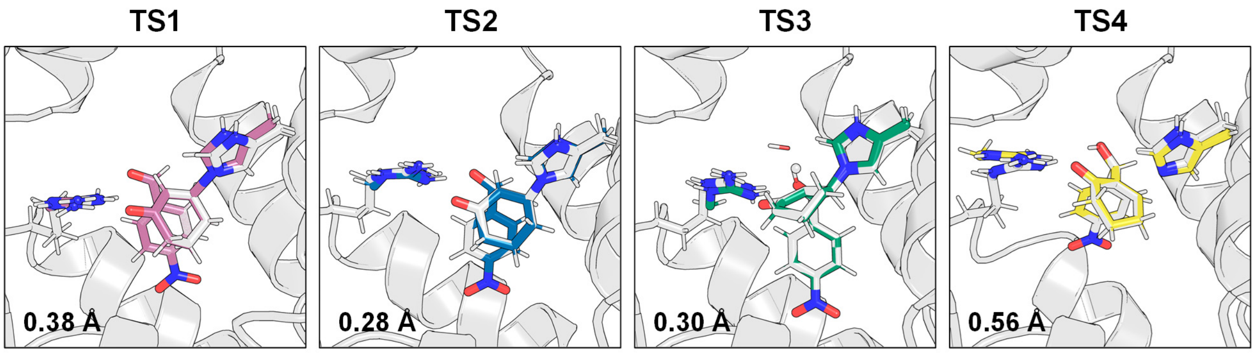


**Supplementary Figure 1. Accurate placement of theozymes on MBHase backbones.** DFT theozyme models (coloured) are superposed with the corresponding theozymes placed by Triad on individual BH32.12 and BH32.8 backbone conformations from their respective ensembles (white). RMSDs between the guanidinium group of Arg124 and the imidazole group of His23 and their corresponding positions in the DFT models are indicated. The close agreement between models highlights the accuracy of theozyme placement. The four theozyme-bound structures shown were subsequently used as inputs for multistate enzyme design.

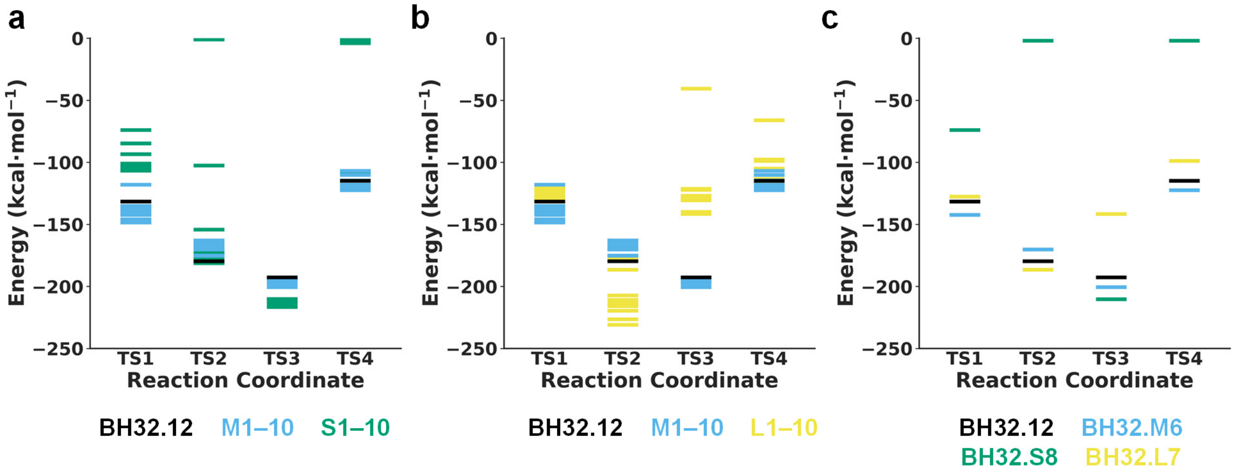


**Supplementary Figure 2. Predicted Triad energy profiles.** (a) Predicted energy profiles of BH32.12 (black), multistate (blue) and single-state (green) designs across the reaction coordinate. Triad-calculated potential energies report the relative favourability of enzyme–transition-state structures but do not represent physical free energies. Multistate designs (M1–10) maintain favourable energies across all transition states, whereas single-state designs (S1–10) preferentially stabilize TS3 at the expense of other transition states. TS2 and TS4 are strongly destabilized in several single-state designs, yielding positive energies capped at 0 kcal $mol^{-1}$ for visualization. (b) Predicted energy profiles of BH32.12 (black), multistate (blue) and LigandMPNN (yellow) designs show that LigandMPNN designs preferentially stabilize TS2 while destabilizing TS3, resulting in greater energetic imbalance than the physics-based multistate designs. (c) Predicted energy profiles of BH32.12 (black), BH32.M6 (blue), BH32.S8 (green) and BH32.L7 (yellow).

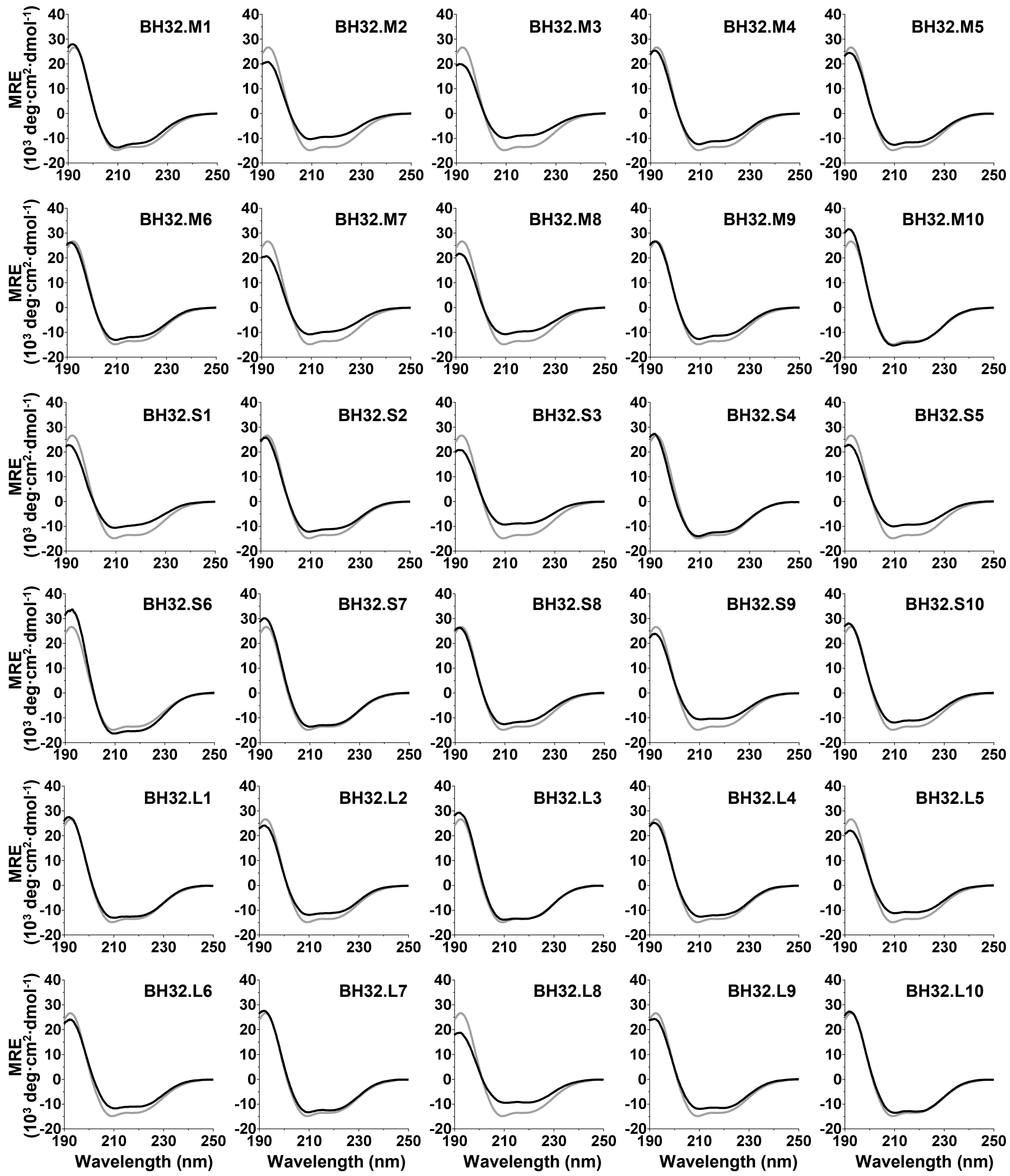


**Supplementary Figure 3. MBHase variants adopt folded structures in solution.** Far-UV circular dichroism spectra of MBHase variants (black) superimposed on the spectrum of BH32.12 (grey).

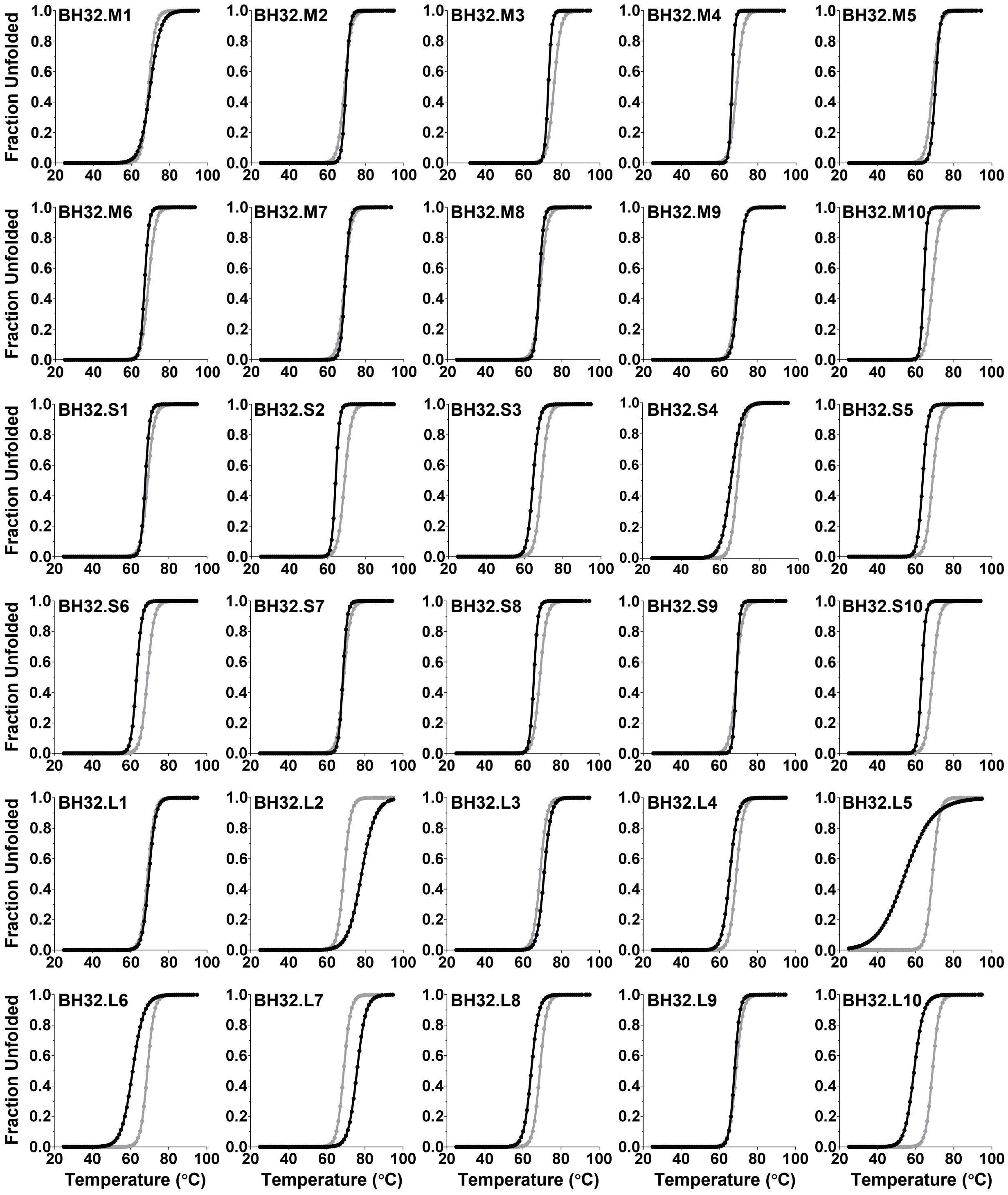


**Supplementary Figure 4. Thermal denaturation of MBHase variants.** Thermal denaturation monitored by circular dichroism at 222 nm indicates that all MBHase variants are stable at room temperature. Data was fit to a two-state unfolding model. Data from the thermal denaturation of BH32.12 is shown in grey for comparison. $T_m$ values are reported in Supplementary Table 6.

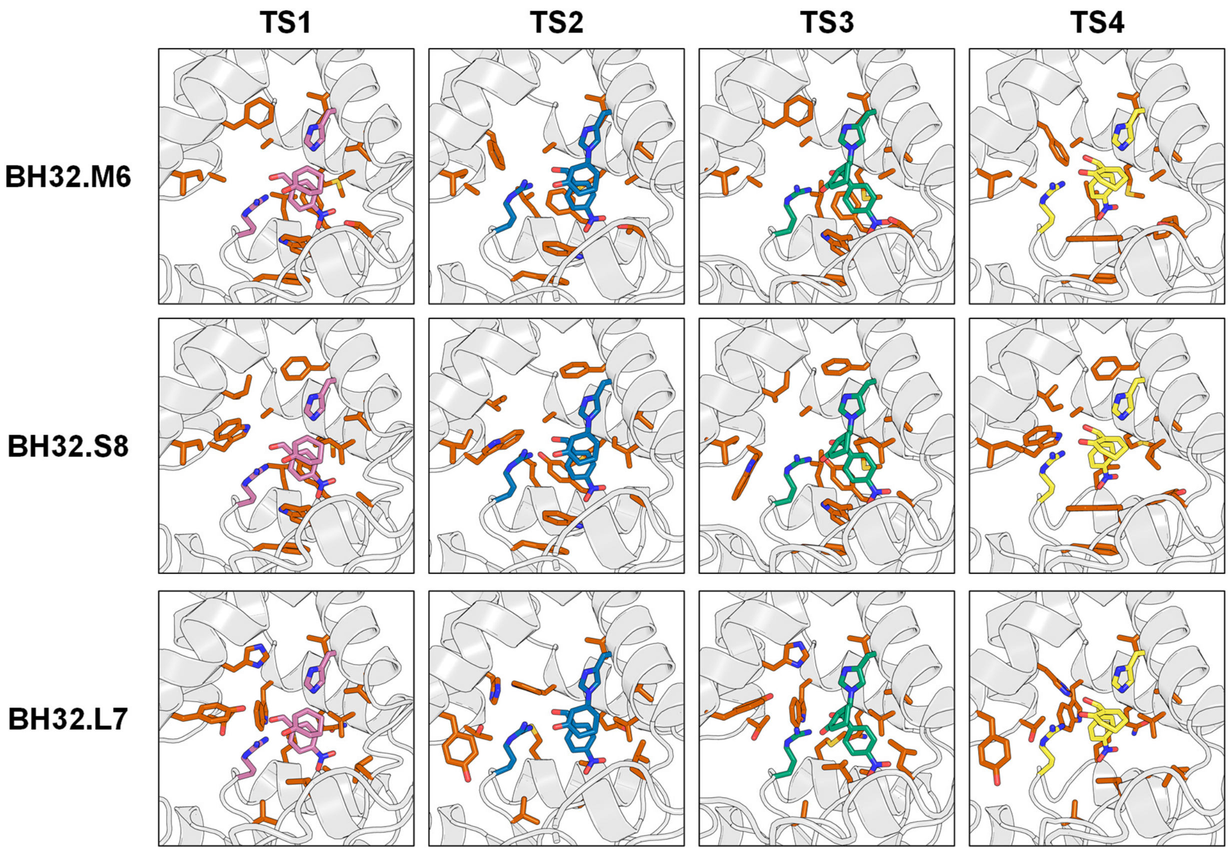


**Supplementary Figure 5. Active-site configurations of designed MBHases.** Residues optimized by Triad during active-site repacking of variants are shown as orange sticks. The catalytic His/Arg dyad and transition states are depicted as pink, blue, green or yellow sticks for TS1, TS2, TS3 or TS4, respectively.

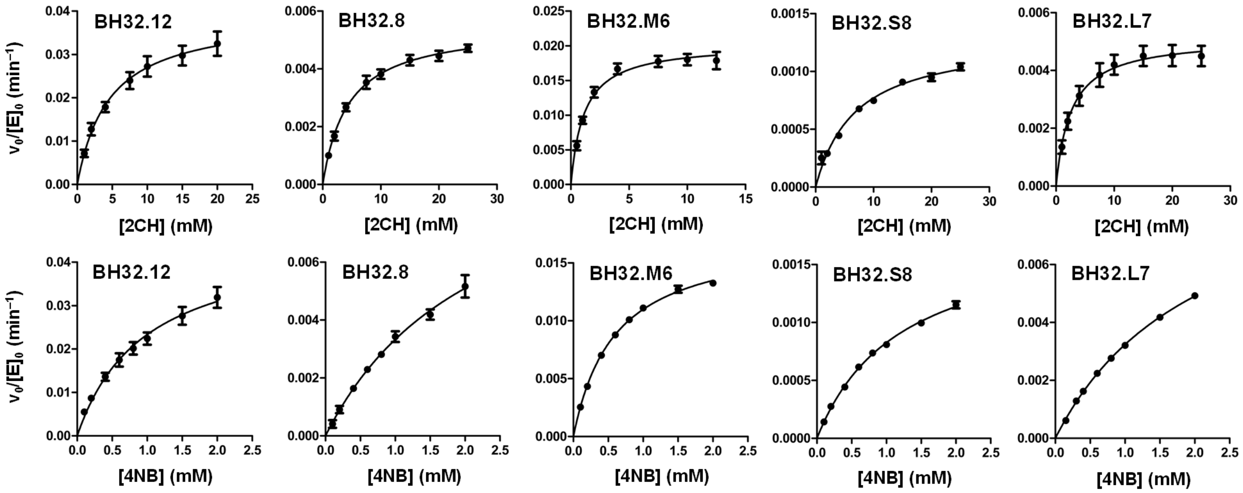


**Supplementary Figure 6. Steady-state kinetics of MBHases.** Michaelis–Menten plots showing normalized initial rates as a function of 2-cyclohexen-1-one (2CH) or 4-nitrobenzaldehyde (4NB) concentration. Data represent the mean of all biological replicates from independent protein batches (n = 8/4, 2/2, 8/3, 6/2, and 5/3 batches for 2CH/4NB measurements of BH32.12, BH32.8, BH32.M6, BH32.S8, and BH32.L7, respectively). Error bars denote s.e.m. Kinetic assays were carried out at 30°C in phosphate buffered saline (pH 7.4) containing 3% acetonitrile (v/v).

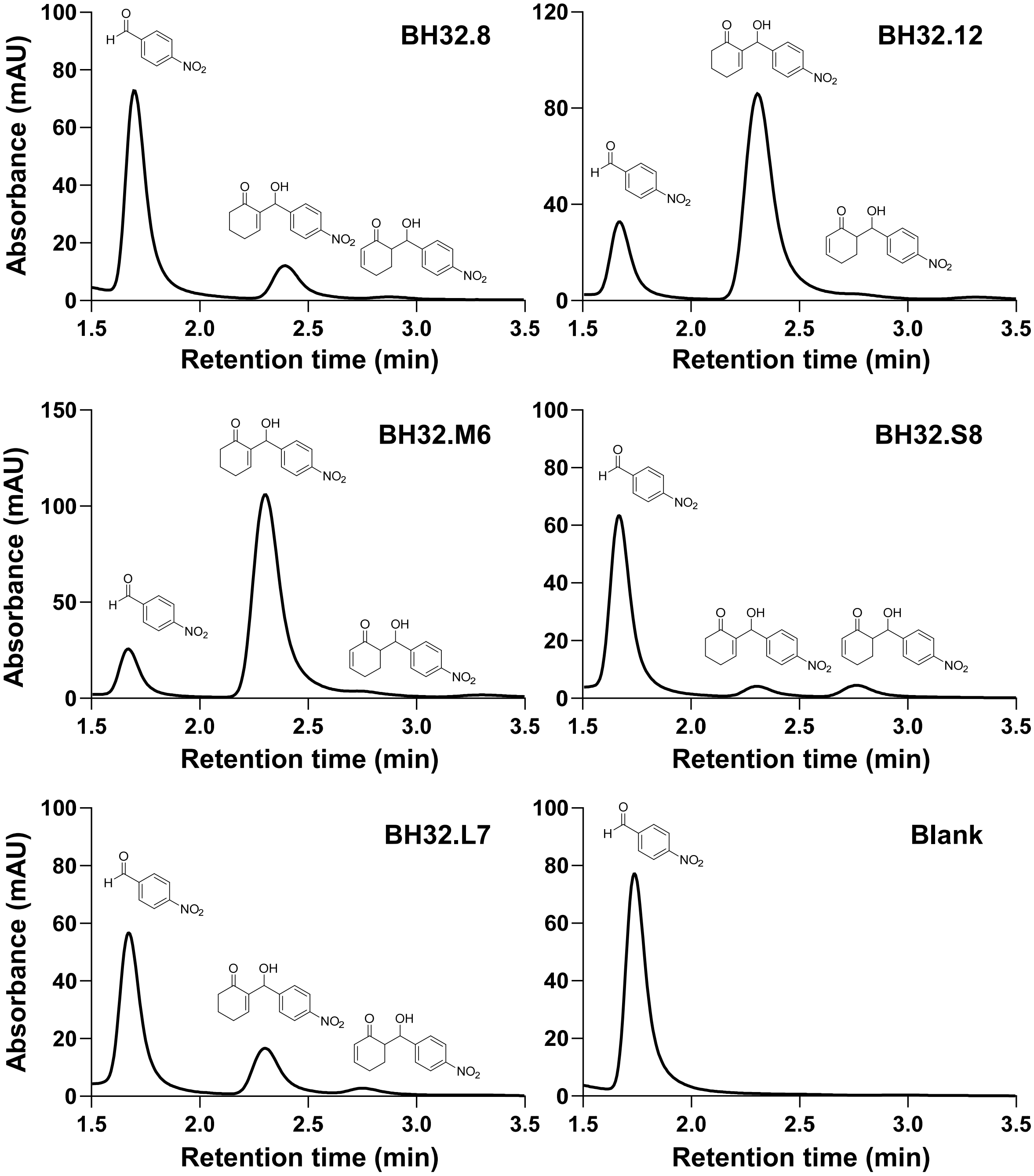


**Supplementary Figure 7. HPLC analysis of MBHase reactions.** Achiral HPLC trace showing the remaining 4-nitrobenzaldehyde substrate and the MBH product and aldol by-product formed after 22 h of incubation. Enzyme reactions were performed with 3 mM 2-cyclohexen-1-one, 0.6 mM 4-nitrobenzaldehyde and 20 µM MBHase (3.3 mol%) at 30 °C in phosphate-buffered saline (pH 7.4) containing 3% (v/v) acetonitrile. The blank contained no enzyme.

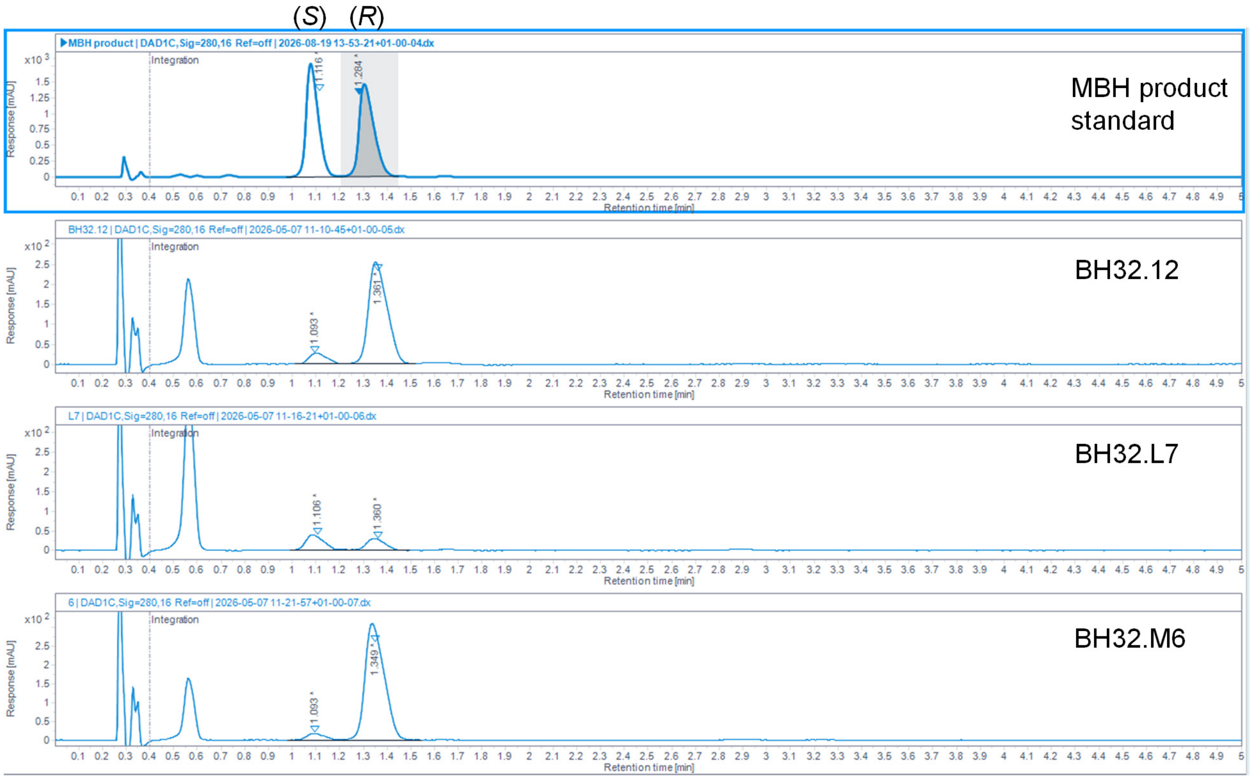


**Supplementary Figure 8. HPLC analysis of MBHase enantioselectivity.** Chiral HPLC trace showing separation of the *R* and *S* enantiomers of the MBH product after 24 h of reaction. Analytical scale enzyme reactions were performed using 3 mM 2-cyclohexen-1-one, 0.6 mM 4-nitrobenzaldehyde and 20 μM MBHase (3.33 mol%) at 30 °C in phosphate-buffered saline (pH 7.4) containing 3% (v/v) acetonitrile.

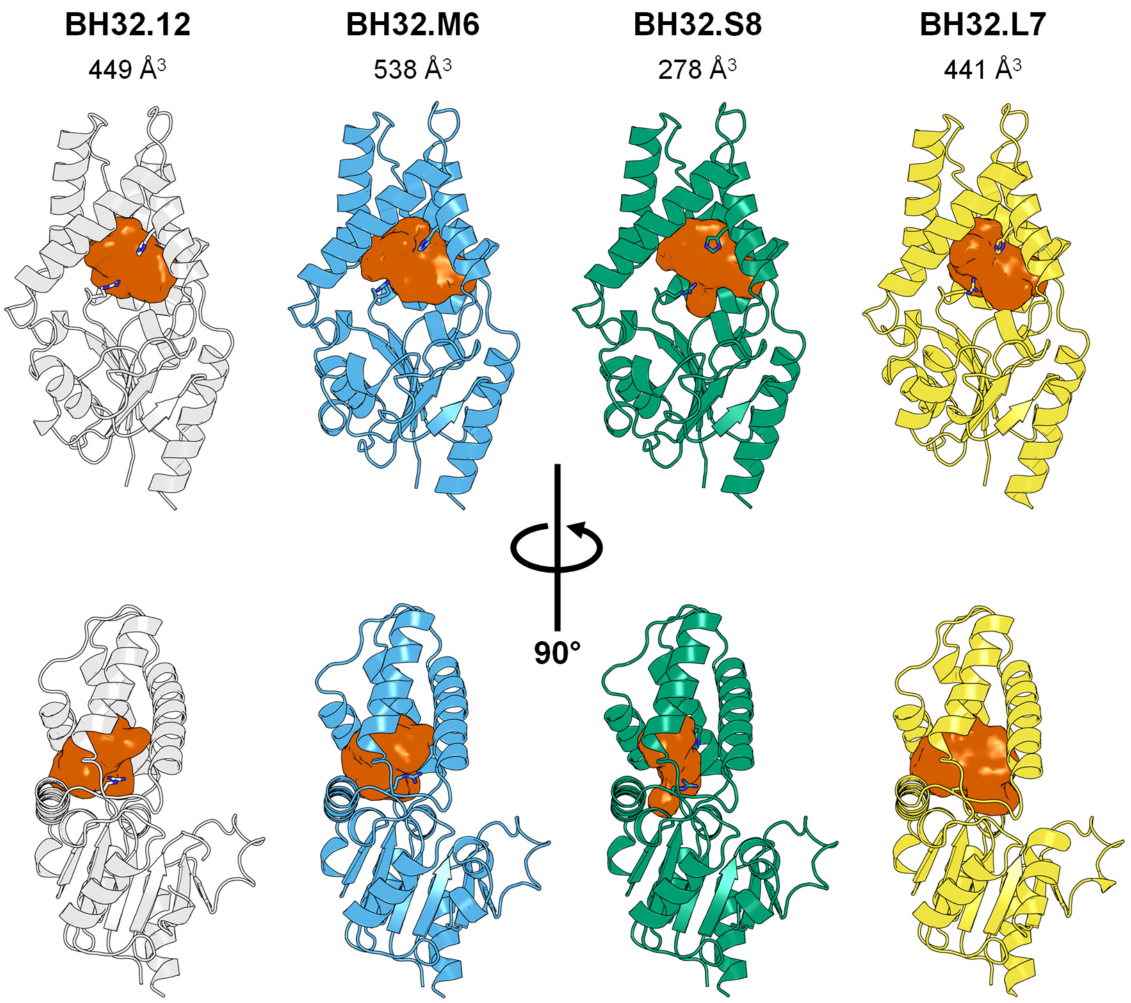


**Supplementary Figure 9. Active-site cavities in MBHase variants.** Cavities (orange) are shown for each MBHase crystal structure, with catalytic residues displayed as sticks. Cavity volumes were calculated using KVFinder on the major conformer of each crystal structure, centering the search region on the TS1 position from the corresponding design models and applying default KVFinder settings.

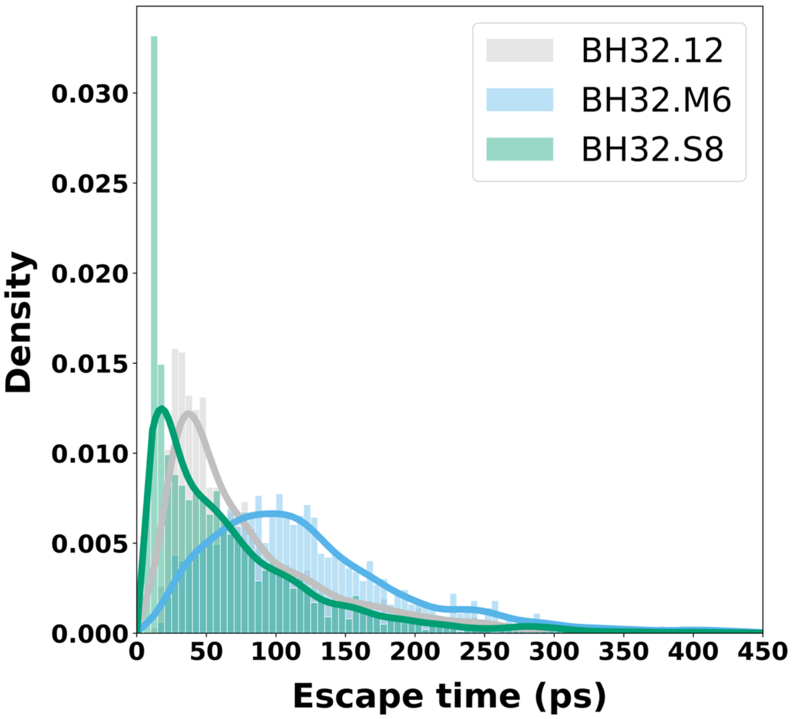


**Supplementary Figure 10. Product release propensity of MBHase variants.** Random-acceleration molecular dynamics (RAMD) were used to compare product-release propensity. In RAMD, a small random force is applied to the product molecule to accelerate its escape from the active site, while the protein and solvent respond dynamically. The resulting escape-time distributions report how readily each MBHase variant releases product, with shorter escape times indicating greater release propensity. See Methods for details.

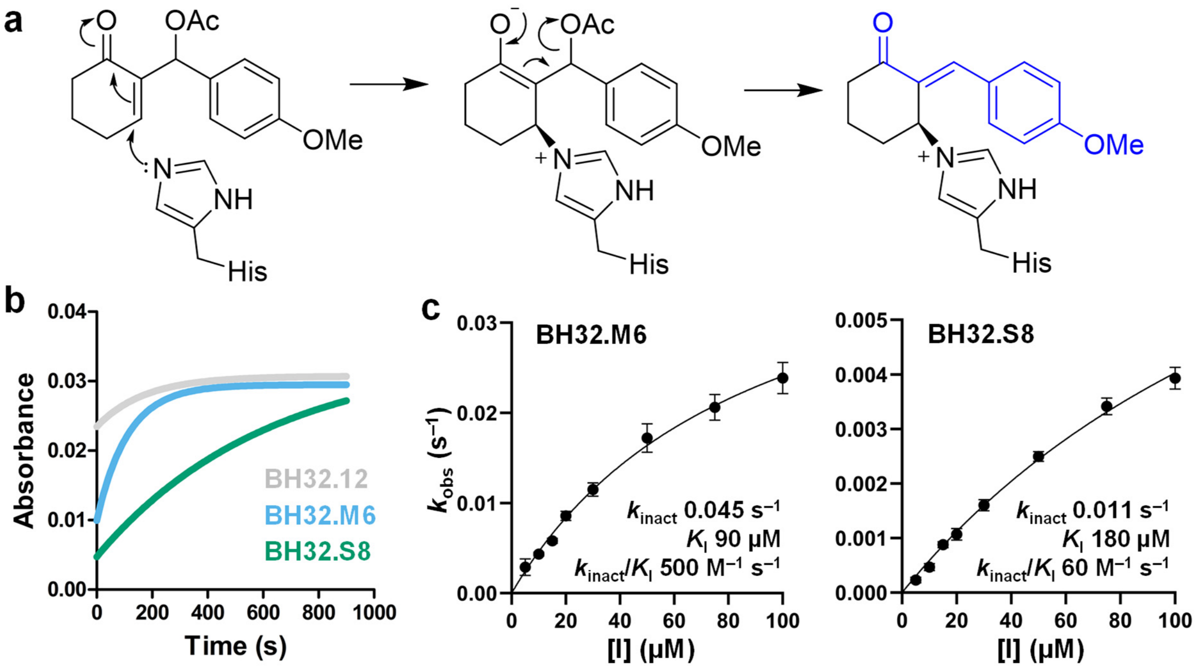


**Supplementary Figure 11. Mechanistic inhibition of MBHases.** (a) Mechanism of MBHase inhibition by the mechanistic inhibitor. After formation of a non-covalent enzyme–inhibitor (EI) complex, the inhibitor undergoes Michael addition by His23, followed by acetate elimination to generate a conjugated π system (blue), which can be monitored by the increase in absorbance at 325 nm. (b) Time course of MBHase variant inhibition. Covalent modification was monitored spectrophotometrically at 325 nm. Reactions contained 4 μM enzyme and 20 μM inhibitor in phosphate-buffered saline (pH 7.4) containing 3% (v/v) acetonitrile at 25 °C. (c) Inhibitor concentration-dependent $k_{obs}$ values were used to determine the inactivation rate constant ($k_{inact}$), inhibition dissociation constant ($K_I$) and inactivation efficiency ($k_{inact}/K_I$) for BH32.M6 and BH32.S8. These parameters could not be determined for BH32.12 because the reaction was too fast to accurately measure $k_{obs}$. For BH32.M6, two independent protein batches were analyzed with four and six technical replicates, respectively; for BH32.S8, two independent protein batches were analyzed with six technical replicates each (mean ± s.e.m.). Here, $k_{inact}$ is the first-order rate constant for covalent modification following EI complex formation, whereas $k_{inact}/K_I$ is the second-order efficiency constant for irreversible inhibition, accounting for both EI complex formation and subsequent covalent modification.

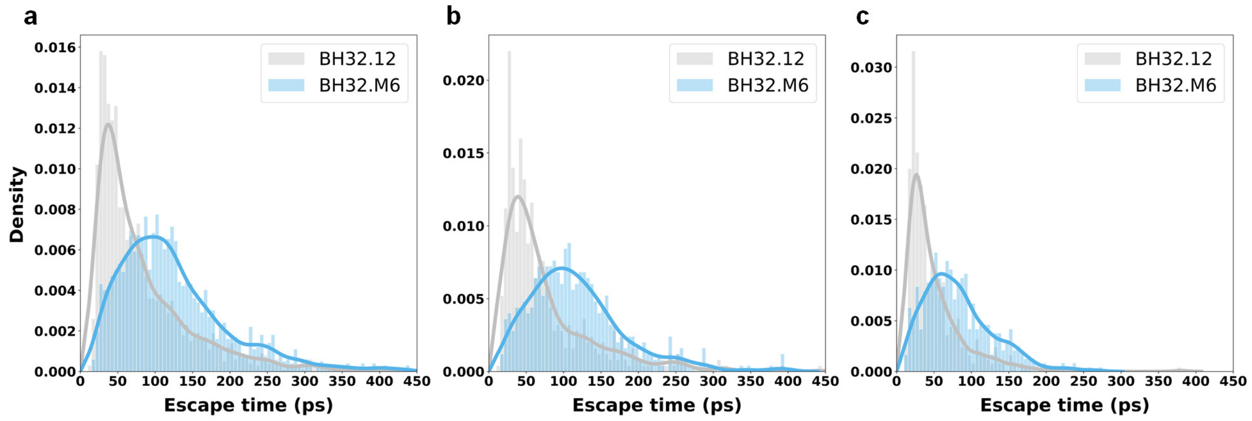


**Supplementary Figure 12. Effect of force F and number of RAMD simulations N on escape time distributions.** (a) F = 450 kJ $mol^{-1}$ $Å^{-1}$, N = 2000; (b) F = 450 kJ $mol^{-1}$ $Å^{-1}$, N = 500; (c) F = 500 kJ $mol^{-1}$ $Å^{-1}$, N = 500.
